# Transcription-based identification of uncharacterized genes in the human immune response

**DOI:** 10.1101/2025.04.22.646545

**Authors:** Emil E. Vorsteveld, Simone Kersten, Charlotte Kaffa, Annet Simons, Peter A.C. ’t Hoen, Mihai G. Netea, Alexander Hoischen

## Abstract

Host-pathogen interactions are shaped by the nature of the pathogen and by host-related factors. The human host responses can be characterized in microbe-stimulated immune cells using transcriptomics. We set out to characterize gene expression changes as a result of microbial *in vitro* stimulation in RPMI-cultured human primary immune cells, using four pathogen ligands representing bacteria (LPS and *S. aureus*), viruses (Poly(I:C)) and fungi (*C. albicans*) for 4 and 24 hours, resulting in a total of 52 samples for analysis. We analyze the transcriptional changes on gene- and pathway level, highlighting common and distinct effects of pathogen stimulation. We highlight genes without a known function that were differentially expressed as a common effect of pathogen stimulation. We find uncharacterized genes such as *KIAA0040* and *CYRIA* to be co-expressed with genes involved in innate immunity and downstream signaling from the PRRs, respectively. We identify 901 variants in uncharacterized genes in a cohort of patients with rare immune disorders, prioritizing 5 candidate variants in 4 genes that could underly these diseases. Our results therefore indicate the potential role of these genes the human immune response and provides candidate variants with potential relevance in rare immune-mediated diseases.

## Introduction

Monogenic defects in genes involved in host response to pathogens underlie inborn errors of immunity (IEI), a group of disorders of the immune system that often remains genetically undiagnosed (1,2). We were motivated by the apparent knowledge gaps in the genes responsible for these rare diseases and other immune-mediated diseases to investigate the transcriptional changes in human immune cells during *in vitro* pathogen stimulation to gain insight into the core genes involved in host-pathogen interactions.

*Ex vivo* stimulation of primary immune cells with pathogens or microbial components are a widely used method for the investigation of host-pathogen interactions, as it mimics *in vivo* pathogen exposure. Stimuli that are commonly used include molecules that stimulate specific pathogen recognition receptors (PRR), such as TLR4 ligand lipopolysaccharide (LPS), Poly(I:C), which stimulates the dsRNA sensor TLR3 and Imidazoquilines, that stimulate TLR7 and TLR8. Additionally, live or heat-killed pathogens can be used in stimulation assays, which simultaneously stimulate a broader range of PRRs. These for instance include the TLRs, NOD- like- and C-type lectin receptors for pathogens such as *S. aureus* or *C. albicans* (3,4).

Transcriptomics has been fundamental in the understanding of various rare disease, including immune-mediated diseases such as IEI. RNA sequencing-based methods (RNAseq) provide diagnostic utility in cases where the presence of deleterious genetic variation influences gene expression or splicing, indicating genetic variants responsible for a patient’s phenotype (5,6). Other efforts into the identification of novel candidate genes for rare disease have for instance exploited gene networks to generate candidate genes or to investigate the relationship between genetic disorders on a molecular level (7–10).

Generating candidate genes for investigation in rare disease cohorts is highly dependent on functional annotations of genes, meaning that genes without a described function remain out of focus in these efforts due to their lack of associations with relevant biological processes (11,12).

Here, we applied bulk 3’ mRNA sequencing (QuantSeq) on *in vitro/ex vivo* microbe-stimulated peripheral blood mononuclear cells (PBMCs) from five healthy individuals (13). PBMCs were stimulated with LPS, *S. aureus*, Poly(I:C) or *C. albicans* for 4 and 24 hours to mimic early- and late innate immune responses to Gram-negative and Gram-positive bacteria, viruses and fungi, respectively. Through differential gene expression analysis, we identify a core set of genes with (candidate) functions in immunity that are differentially expressed as a result of *in vitro* microbe stimulation. We highlight genes without a known function that are co-expressed with modules of known immune genes, which may imply these to play a role in core functions in the human immune response and may further implicate these genes as novel candidates for immune-mediated diseases such as IEI, which we investigate through a cohort of IEI patients that underwent exome sequencing.

## Methods

### Ex vivo immune cell stimulation

Isolation of peripheral blood mononuclear cells (PBMCs) was conducted as described elsewhere (14,15). In brief, PBMCs were obtained from blood by differential density centrifugation over Ficoll gradient (Cytiva, Ficoll-Paque Plus, Sigma-Aldrich) after 1:1 dilution in PBS. Cells were washed twice in saline and re-suspended in serum-free cell culture medium (Roswell Park Memorial Institute (RPMI) 1640, Gibco) supplemented with 50 mg/mL gentamicin, 2 mM L-glutamine and 1 mM pyruvate. Cells were counted using a particle counter (Beckmann Coulter, Woerden, The Netherlands) after which the concentration was adjusted to 5 × 10^6^/mL. *Ex vivo* PBMC stimulations were performed with 5×10^5^ cells per well in round-bottom 96-well plates (Greiner Bio-One, Kremsmünster, Austria) for 24 hours at 37°C and 5% carbon dioxide. Cells were stimulated with lipopolysaccharide (*E. coli* LPS, 10 ng/mL), heat-killed *Staphylococcus aureus* (ATCC25923, 1×10^6^/mL), TLR3 ligand Poly I:C (10 µg/mL), heat-killed *Candida albicans* yeast (UC820, 1×10^6^/mL), or left untreated in regular RPMI medium as normal control. After the incubation period of 24h and centrifugation, supernatants were collected and stored at -80°C until further processing. For the RNA isolation, cells were stored in 350 µL RNeasy Lysis Buffer (Qiagen, Rneasy Mini Kit, Cat nr. 74104) at −80°C until further processing.

### QuantSeq 3’mRNA sequencing

RNA input was normalized to 200 ng for all samples/donors. Libraries were generated using the QuantSeq 3’ mRNA-Seq Library Prep Kit-FWD (Lexogen) in accordance with the manufacturers’ protocol (13). In order to ensure high quality libraries, two separate preparations were performed, limiting the number of samples to 30 per preparation. End-point PCR was performed with 19 – 22 cycles, as indicated by a quantitative PCR on a 1:10 aliquot of a subset of double stranded cDNA libraries. Accurate quantification and quality assessment of the generated libraries was performed using Qubit dsDNA High Sensitivity assay (Thermo Fisher Scientific) and Agilent 2200 TapeStation (High Sensitivity D1000 ScreenTape, Agilent). Molarity of individual libraries was calculated using the cDNA concentration (Qubit) and average fragment size (TapeStation). Safeguarding sufficient read depth for each sample, libraries were split in two separate runs. In each run, the baseline RPMI condition across all donors and time-points was included, in turn allowing sequencing bias assessment. Per pool, a total of 35 samples/cDNA libraries were pooled equimolar to 100 fmol. After a final dilution of both pools to a concentration of 4 nM, they were sequenced on a NextSeq 500 instrument (Illumina) with a final loading concentration of 1.4 pM.

### Quantification and quality control

The quality of the obtained sequencing data was assessed using FastQC (v0.11.5, Babraham Bioinformatics), followed by removal of adapter sequences and poly(A) tails using Trim Galore! (v0.4.4_dev, Babraham Bioinformatics) and Cutadapt (v1.18) (16). Filtered and trimmed reads were mapped to the human reference genome (hg38/GRCh38.p12) using the STAR aligner (v2.6.0a) (17) and subsequently counted using HTSeq-count (v0.11.0) (18). We excluded samples with <1.5*10^6^ assigned reads for further analysis (n=6), leaving 52 samples for analysis (Table S1-2). These data were previously used in the validation of expression data from long read transcriptomics generated from five samples from one of the five individuals included in the present study (19,20).

Given our primary interest in protein coding genes, we removed genes labeled as mtrRNA, mttRNA, snRNA, snoRNA, miscRNA and lincRNA, as registered in Ensembl v95 from our count matrix. We converted each Ensembl gene ID to gene symbols using the same version of the database. We retained genes with ≥10 total reads across all samples.

### Gene body coverage

We assessed the sequencing coverage across the gene body using the RESeQCoverage.py script from the RSeQC Python package (v5.0.0) using the default set of 3802 housekeeping genes (https://sourceforge.net/projects/rseqc/files/BED/Human_Homo_sapiens/hg38.HouseKeepi ngGenes.bed.gz/) (21,22). We scaled the coverage per percentile using the rescale() function from the scales R package (v1.2.1) (23). The coverage per percentile was plotted per sample. As expected, we found high sequencing coverage at the 3’ ends of transcripts and low coverage at the 5’ ends of transcripts (Figure S1).

### Gene expression correlation

We assessed the correlation of the RPMI samples to control for sequencing bias between the samples. We generated Pearson correlations between biological and technical replicates using the cor() R function. We find a correlation between technical replicates (baseline RPMI), i.e., individual cDNA libraries included in the separate sequencing runs, indicating limited sequencing bias within our experiments (Pearson R2=1 for all samples, Figure S2A-B). Similarly, the Pearson correlation (R2) of all biological replicates was 0.97, suggesting minimal variation in our dataset was introduced during the experimental or library preparation procedures (Figure S2C-D).

### Principal component analysis

To gain insight into the relationship between the samples in our dataset, we performed a principal component analysis (PCA) on the DESeq2-normalized gene counts (24). We used the plotPCA() function from the DESeq2 R package (v1.40.2) to generate the principal components from the normalized expression data, using the default top 500 genes with the highest variability (24). We applied PCA to the full dataset and to the 4 and 24 hour data separately.

### Hierarchical clustering

We performed k-means clustering to find similar samples from the normalized expression data, generated using the rlog() function from the DESeq2 R package (v1.40.2) (24). We selected the top 500 genes with the most variation and determined the optimal number of clusters using the fviz_nbclust() function from the factoextra R package (v1.0.7) using seed #24 (25). We found an optimal number of 6 gene clusters and 6 sample clusters (Figure S3, Table S3). The clustering was visualized using the pheatmap() function from the pheatmap R package (v1.0.12) (26).

### Gene set co-regulation

In order to find gene sets that were correlated, we applied a gene set co-regulation analysis (GESECA). Genes with at total count ≥10 across all samples were included. As suggested by the authors, we normalized the expression levels using the normalizeBetweenArrays() function from the limma R package (v3.56.2) (27) and ran GESECA with seed #1 using the geseca() function from the fgsea R package (v1.26.0) (28). We used Gene Ontology Biological Process gene sets from the Molecular Signatures Database (MSigDB) with between 100 and 500 genes to find correlations of these gene sets across samples (29,30). Gene sets with a Benjamini-Hochberg adjusted P value <0.01 were considered significant (Table S4).

### Gene set enrichment analysis

We assessed pathway enrichment using gene set enrichment analysis (GSEA) (31), as implemented in the fgsea R package (v1.26.0) (28). We used the fgsea() function from this package with the DEGs generated for each stimulus-time point combination. We used the Biological Process gene sets from Gene Ontology from the Molecular Signatures Database (MSigDB) with between 100 and 500 genes (Table S5) (29,30). We similarly used genes specifically expressed in 15 immune cell types for GSEA from DICE (100 genes each) to infer cell-type specific transcriptional changes (Table S6) (32). Gene sets with a Benjamini-Hochberg adjusted P value <0.01 were considered significantly enriched.

To gain insight into the functional overlap between stimulus conditions, we gathered a representative subset of the overlapping pathways that were upregulated at both time points in all conditions using REVIGO (33). We supplied the Gene Ontology IDs of the gene sets that were enriched for all stimuli at both time points (N=108) to REVIGO and used the default settings to generate a summary pathway set (Table S7). We visualized the “Interactive Graph” output in Cytoscape (v3.10.0) (34) (File S1). Additionally, we obtained the semantic similarity (SimRel) matrix generated for the set of pathways by REVIGO and classified these using kmeans clustering. We determined the optimal number of clusters using the fviz_nbclust() function from the factoextra R package (v1.0.7) (25). This yielded an optimal number of 5 clusters (Figure S4). We used the pheatmap() function from the pheatmap R package (v1.0.12) to visualize the pathway clusters (26).

### Differential gene expression

In line with our aim to gain a deeper understanding of the general pathogen-induced host response, PBMC donors were considered biological replicates and compared to the control condition (RPMI). Differential gene expression analysis was carried out using DESeq2 (v1.40.2) (24), including logFold Shrinkage with apeglm (v1.22.1) (35), as described by the authors. Genes with an FDR-adjusted P value <0.01 and |^2^logFC| ≥ 1 were considered differentially expressed genes (DEG). As a quality control measure, we visualized the DEGs in a volcano plot per stimulus and time point combination (Figure S5, Table S8).

We visualized the DEGs associated with KEGG “Toll-like receptor signaling pathway” system (hsa04620) (36). We used the Omics Visualizer plugin in Cytoscape to produce a donut diagram for each time point associated with the log2 fold-changes generated from DESeq2 (34,37).

### Gene expression validations using external datasets

For validation of our gene counts, we used external datasets of pathogen stimulated PBMCs for which the processed counts files were available in the Gene Expression Omnibus (38).

We used data from a dataset of LPS-stimulated PBMCs (39), Poly(I:C)-stimulated PBMCs (40), a dataset of PBMC stimulated with either *C. albicans* and *C. auris* from Bruno et al. (41) and a dataset of PBMCs stimulated with *C. albicans*, *A. fumigatus* or *R. oryzae* (42). No suitable datasets using *S. aureus* stimulation experiments were identified. We selected samples with matching stimulation times for each dataset, merging the gene counts of both datasets by Ensembl ID and corrected for dataset-specific effects using the *removeBatchEffect()* function from the limma R package (v3.56.2) (27). We performed principal component analysis and DEG generation for each dataset following the same procedure described above. We used a Fisher’s exact to assess the overlap between our DEGs and those in the external datasets.

We assessed the correlation between the log2 fold-change values between DEGs of the matching conditions and time points with our dataset with a linear regression using the lm() function in R (Table S9).

### Functional classification of DEGs

Providing an additional layer of biological and molecular interpretation, we classified the DEGs for their association to the immune system and function in immune related processes. Genes were designated to one of four classes: i) established inborn errors of immunity (IEI) disease genes (n=482), as defined by the International Union of Immunological Societies (IUIS) 2020 (1) and our *in silico* diagnostic PID immune panel (DG 3.3.0, Genome Diagnostics Nijmegen) (43); ii) genes with known immune association (n=1909, excl. known IEI), as defined by Immunogenetic Related Information Source (44), InnateDB (45) and Reactome’s Immune system pathway (46); iii) uncharacterized genes, i.e. genes with a UniProt annotation score of ≤2 (generated by extracting all entries by annotation score) (47,48); and iv) other genes, composing all other differentially expressed genes (Table S10).

To assess our classification of uncharacterized genes, we gathered the number of publications associated with each gene in our dataset using the Find My Understudied Genes database (FMUG) (11). We were able to identify the number of publications for 29810 genes in our dataset (64.8%; Figure S6A, Table S11), which includes 19,206 protein coding genes (96.8%; 19,844 in total). We identified 3320 DEGs in this dataset (92.8%, Figure S6B, Table S11). We compared the number of publications between characterized and uncharacterized genes using a Wilcoxon Signed Rank test.

### Functional enrichment analysis of core DEGs

We assessed the functional enrichment of the overlapping DEGs between conditions at either time point using g:Profiler as implemented in the gprofiler2 R package (v0.2.3) (49,50). Bonferroni-corrected P values <0.01 were considered significant (Table S12).

We further assessed the enrichment of IEI- and immune-related genes among the core DEGs, using the remaining DEGs as background. We tested for enrichment of genes in our in-house diagnostic IEI gene panel (DG 3.3.0, Genome Diagnostics Nijmegen) (43), genes in IUIS table 6 (defects in intrinsic and innate immunity N=65), genes with known immune association (N=1909, excl. known IEI genes), as defined by Immunogenetic Related Information Source (44), InnateDB (45) and Reactome’s Immune system pathway (46). We further tested for enrichment of mouse knockouts with immune-related phenotypes in the JAX Mouse Genome Informatics resource (retrieved from MGI). We defined immune-related phenotypes to include all genes that include the “immu” string in the high-level or low-level phenotype descriptions in MGI (51). We used a Fisher’s exact test. We performed an enrichment analysis with a Fisher’s exact test and considered Bonferroni-corrected P values <0.01 as significant (Table S13).

### GWAS hits associated with pathogen response DEGs

We downloaded the GWAS signals registered in the GWAS catalog (v1.0.2) (52). We gathered the genes associated with each of the 10,055 phenotypes by their phenotype mapping identifier (EFO). This yielded an average of 18.4 genes per trait (range 0-6530). As a high number of traits are associated with a small number of genes (N=8122 with N≤10, 80.8%), we therefore opted to include only traits with >10 associated genes (N=1933, 19.2%, Figure S7A). We calculated the enrichment of all GWAS gene sets among the DEGs identified in each condition-time point combination using a Fisher’s exact. We considered gene sets with a Bonferroni-corrected P value <0.01 to be significantly enriched (Table S14).

### eQLTs associated with pathogen response DEGs

We analyzed the eQTLs identified using publicly available expression data from 15 immune cell types (32). The genes mapped to the eQTLs (eGenes) are generally distinct between cell types, with between 452 and 955 unique genes per cell type, with 452 genes shared by all cell types (Figure S8A). We assessed the enrichment of eGenes among the DEGs identified at either time point for each of the four stimulus conditions using a Fisher’s exact. We considered gene sets with a Bonferroni-corrected P value <0.01 to be significantly enriched (Table S15).

### Gene network analysis of core DEGs

We analyzed the DEGs expressed in response to all four stimuli at 4 and 24 hours separately through a gene network analysis. The gene names were used to generate a network in Cytoscape (v3.10.0) using STRING data (v12.0) (34,53). The largest subnetwork was chosen, and a functional enrichment was performed using gProfiler implemented in the gprofiler2 R package (v0.0.2) (49,50). We selected the Gene Ontology Biological Process gene sets and built an enrichment map with a connectivity cutoff (Jaccard similarity) of 0.4 using Cytoscape’s EnrichmentMap plugin (v3.3.0) (30). We selected the largest subnetwork and manually annotated sets of connected pathways by their general biological function (Table S16-19).

### Co-expression analysis

We applied a weighted gene co-expression network analysis (WGCNA) using the WGCNA R package (v1.72.1) (54). We generated a sample tree using the hclust() R function. We found the samples were separated by time (Figure S9A). We therefore opted to generate two separate analyses for each time point (Figure S9B-C). As suggested in the WGCNA manual, we selected a soft threshold for network generation with a scale free topology model fit of ≥0.8, while maximizing the mean connectivity. This resulted in a soft cutoff of 5 for the 4 hour data (Figure S9D-E) and 10 for the 24 hour data (Figure S9F-G). We constructed the signed networks using the blockwiseModules() function with the matching soft cutoff value, a height of 0.25, a deep split level of 2, a maximum block size of 5000 genes, a minimum block size of 30 genes and seed #1234. The topological overlap matrices were generated using the TOMsimilarityFromExpr() function. Hierarchical clustering of genes was based on the TOM dissimilarity measure (1-TOM) (Table S20).

In order to assess whether the gene modules generated using WGCNA are induced by our pathogen stimuli, we assessed their enrichment as described previously using GSEA implemented in the fgsea R package (v1.26.0) (28), using the DEGs from the matching time point for each gene module. We considered enrichments with a Benjamini-Hochberg adjusted p value <0.01 to be significant (Table S21).

We analyzed each gene module for functional enrichment using g:Profiler implemented in the gprofiler2 R package (v0.2.2) using Gene Ontology, Reactome, CORUM, the Human Protein Atlas and KEGG gene sets. For a simplified visualization, we used only Gene Ontology gene sets (30,36,46,49,50,55,56). We considered gene sets with an adjusted P value <0.01 to be significant (Table S22).

We visualized the gene modules by gathering the correlations of the genes in each WGCNA module with the TOMsimilarityFromExpr() function from the WGCNA package for each time point. As the networks differ in size, we selected a correlation cutoff of >0.3 for MEyellow_24hr, and >0.2 for MEgreen_24hr. Subsets of each network were visualized in Cytoscape (v3.10.2) (34).

### Cell-type level expression of uncharacterized genes

We investigated the expression of *KIAA0040* and *FAM49A (CYRIA)* using expression data from single cell RNA sequencing data available on the CELLxGENE platform (57,58). We filtered for cell types in blood that included ≥1000 cells and obtained the scaled expression levels (Table S23). We further investigated the expression in three datasets of bulk sequencing of sorted PBMCs that are included in the Human Protein Atlas (55); 18 cell population from Uhlen et al. (59), 29 cell types from Monaco et al. (60) and 13 cell types from Schmiedel et al. (32) (Table S24).

### Co-expression from published data

We extracted gene-gene correlation data for uncharacterized DEGs *KIAA0040* and *FAM49A (CYRIA)* using the correlationAnalyzeR R package (v.1.0.0), which exploits the large-scale gene expression database ARCHS4 (61,62). We selected datasets defined as “normal” and extracted the Pearson correlation coefficients using the analyzeSingleGenes() function. We performed GSEA using the ranked correlation coefficients with the fgsea() function from the fgsea R package (v1.26.0) (28). We used Gene Ontology Biological Process gene sets with between 100 and 500 genes (30). We considered gene sets with an adjusted P value <0.01 to be significant (Table S25).

### Rare variant analysis

We analyzed rare variants in the uncharacterized genes found to be DE in our study in a large cohort of patients with IEI that underwent whole exome sequencing in their diagnostic procedure (N>1,300) (63).

To ensure the uncharacterized genes we identified in the current study were included in the exome data, we assessed the coverage of MANE transcripts in exome enrichment kits. We downloaded the exome probesets for Agilent SureSelect V4 and V5 kits and for Twist Exome 2.0, as these are the most common in our clinical exome data and generated the coverage of the MANE transcripts of all DEGs we identified (GRCh38.v1.4) using the “coverage” function from the BedTools package (64). The coverage of exons of uncharacterized genes was significantly lower than for all other genes (Wilcoxon test; Agilent V4 P=2.39e-12; Agilent V5 P=3.94e-08; Twist 2.0 P<2.2*10^-16^).

Similar to previously described analyses of rare variants in patients with rare disease, we filtered for variants that are rare or absent in large databases of genetic variation such as gnomAD, dbSNP and in our in-house exomes database (>30,000 patients) (65–68). We separately analyzed variants by effect (missense, canonical splice sites, stops/frameshifts) at <0.1% allele frequency and homozygous variants (≥80% VAF) at <0.1% and 1% AF.

### Data analysis

We used R (v4.4.0) with tidyverse (v2.0.0) packages for data analysis and plotting (69,70). We used the venn() function from the venn R package (v1.12) to plot overlaps between sets of genes and pathways (71).

We developed an interactive app for the visualization, analysis and download of the DEGs and pathway analysis results using the Shiny framework (72), accessible at https://emilvorsteveld.shinyapps.io/app_de/.

## Results

We stimulated PBMCs from five healthy donors with four microbial stimuli; *E. coli* LPS, heat-killed *S. aureus* and *C. albicans* and Poly(I:C), using RPMI-cultured PBMCs as a control.

Stimulations were carried out for 4 or 24 hours as previously described, resulting in 52 samples for analysis (14). We profiled the transcriptome of the PBMCs using QuantSeq, a short-read RNAseq method that provides coverage at the 3’ ends of transcripts, allowing for gene-level quantification of the transcriptome (Figure 1A and S1) (13).

**Figure 1:**
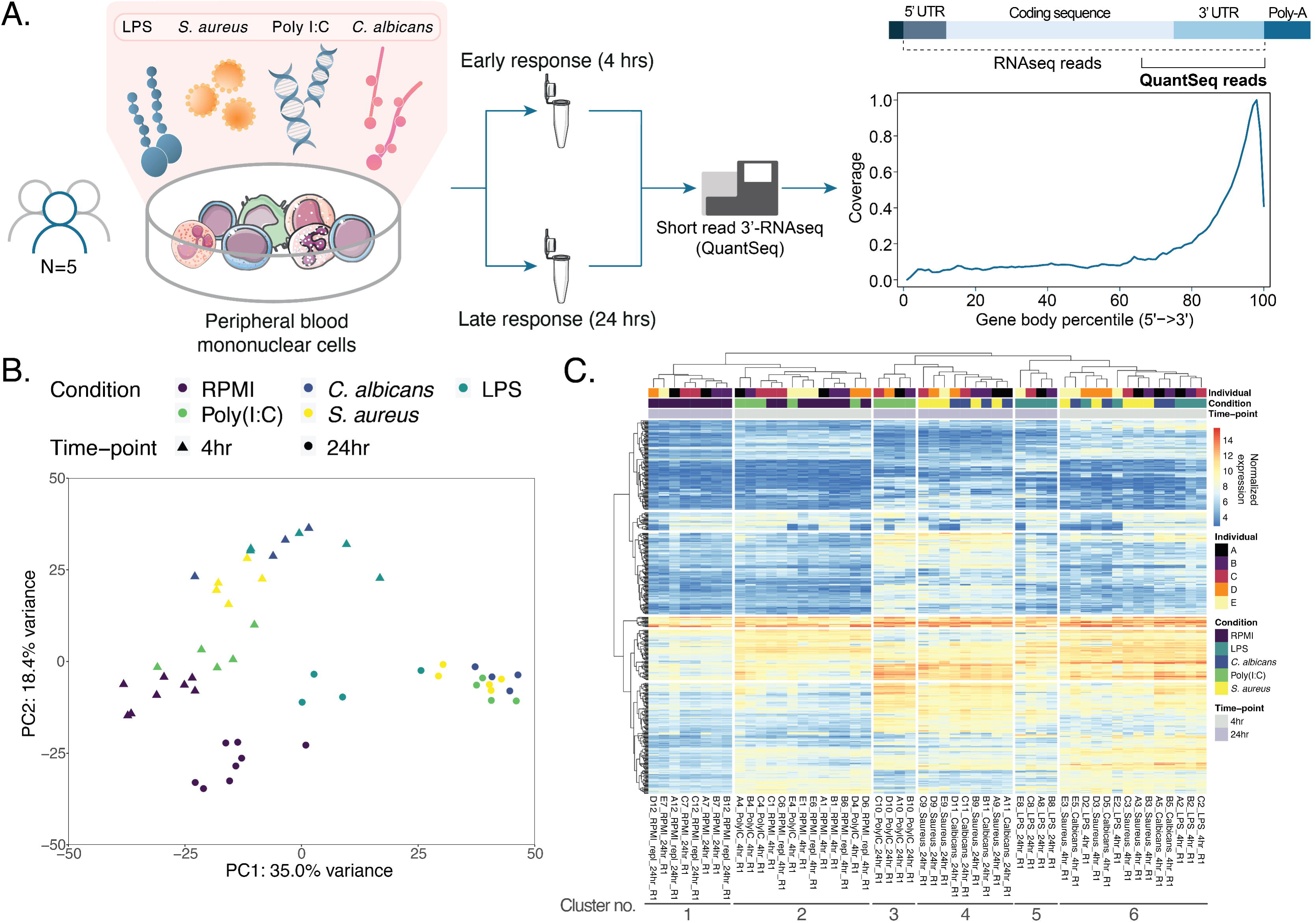
*Workflow and quality control.* A) The experimental workflow of the study. PBMCs of five donors were stimulated with LPS, *S. aureus,* Poly(I:C) or *C. albicans* for 4 and 24 hours. To gain insight into the changes in the transcriptome upon pathogen stimulation, we applied 3’ mRNA sequencing, which provides gene-level quantification of gene expression. B) Principal component analysis of the expression data. Colors indicate conditions and shapes indicate stimulation time. C) Heatmap of clustered samples based on K-means clustering (6 sample clusters and 6 gene clusters; Figure S3).

### Transcriptome-level differences between control- and pathogen-stimulated PBMCs are dependent on stimulus and stimulation time

Using principal component analysis (PCA), we found the first and second principal component (PC) to represent 35.0 and 18.4% of the total variance. We found the samples to be differentiated by condition and time point, represented by a combination of the first two PCs (Figure 1B). Performing PCA separately on the 4- and 24 hour stimulated samples reveals differences between the samples by condition, represented by the first PC (38.0 and 48.5% variance, respectively; Figure S10A-B). We additionally find both stimulated and control samples at the 4 hour time point to separate by sex of the individual donors on the second PC (16.9% variance; male A-C and female D-E), potentially reflecting sex differences in early immune responses (Figure S10A) (73). We did not observe this effect at the 24 hour time point (Figure S10B). The second PC at the 24 hour-time point samples separates the LPS-stimulated samples from all other samples (16.7% variance; Figure S10B). We therefore reasoned that stimulus and time point are responsible for a higher degree of variability between samples than inter-individual differences between donors on gene transcription.

Using a clustering analysis, we find stimulus and control samples at either time point form separate clusters. The control samples at 24 hours form a separate cluster (cluster 1), while the 4 hour control samples cluster together with Poly(I:C)-stimulated samples, reflecting the results from the PCA, further suggesting a less efficient induction of the immune response for this stimulus (Figure 1B-C and S10A-B). We find separate clusters for 24-hour Poly(I:C)- and LPS-stimulated samples (clusters 3 and 5; Figure 1C), while *S. aureus-* and *C. albicans*-stimulated samples form a cluster together at this time point (cluster 4; Figure 1C). Samples stimulated for 4-hours with LPS, *S. aureus* or *C. albicans* also cluster together (cluster 6; Figure 1C).

### Gene co-regulation analysis highlights gene sets with correlated expression patterns in stimulus- and control conditions

In order to further characterize the transcriptional differences between our samples, we performed a gene set co-regulation analysis (GESECA), which allows for the identification of gene sets with correlated expression patterns across samples in multi-conditional data (28). We find gene sets with functions such as cytokine responses and viral and bacterial responses to have high transcriptional correlation across conditions, with high expression in all microbe-stimulated conditions and low expression in control samples. In contrast, genes involved in cytoplasmic translation showed high expression in controls and low expression in stimulus conditions (Figure S11A-B, Table S4), suggesting translational inhibition upon pathogen stimulation, likely reflecting previous findings of a downregulation of these processes in the immune response to various pathogens (74–76).

### Pathway-level analysis of the transcriptomic response to in vitro microbial stimulation

We applied gene set enrichment analysis (GSEA) to detect pathways induced by pathogen stimulation at the two stimulation time points, using RPMI-cultured PBMCs as control (Figure 2A-B and S12A, Table S5, Additional file 1). We find a higher number of induced gene sets at the 24 hour time point than at the 4 hour time point (Figure 2A-B and S12A, Table S5 and S26). The former also displayed a higher degree of overlap, especially considering the upregulated gene sets at this time point. Especially gene sets induced by *S. aureus* and *C. albicans* were highly similar at this time point (Figure 2A-B, S12A, Table S5). We find an induction of expected pathways involved in immunity-related functions at both time points, with more diverse pathways induced upon 24 hour stimulation, such as gene sets with functions in metabolism. We find an LPS-specific induction of lipid transport and lipid membrane structural functions, including functions involved in the encapsulation-related processes (Figure 2B, Table S5 and S26). We find a downregulation of gene sets involved in RNA processing, splicing and metabolism, ribosomal function and chromatin remodeling specifically for 24-hour LPS stimulation (Figure 2B and S12B-E, Table S5 and S26). We find a common downregulation of microtubule polymerization and depolymerization shared between LPS, *C. albicans* and *S. aureus* at the 4 hour time point (Figure 2B, Table S5 and S26), potentially reflecting previous findings where LPS stimulation to affects microtubule organization, with effects on cell shape and migration in leukocytes (77,78).

**Figure 2:**
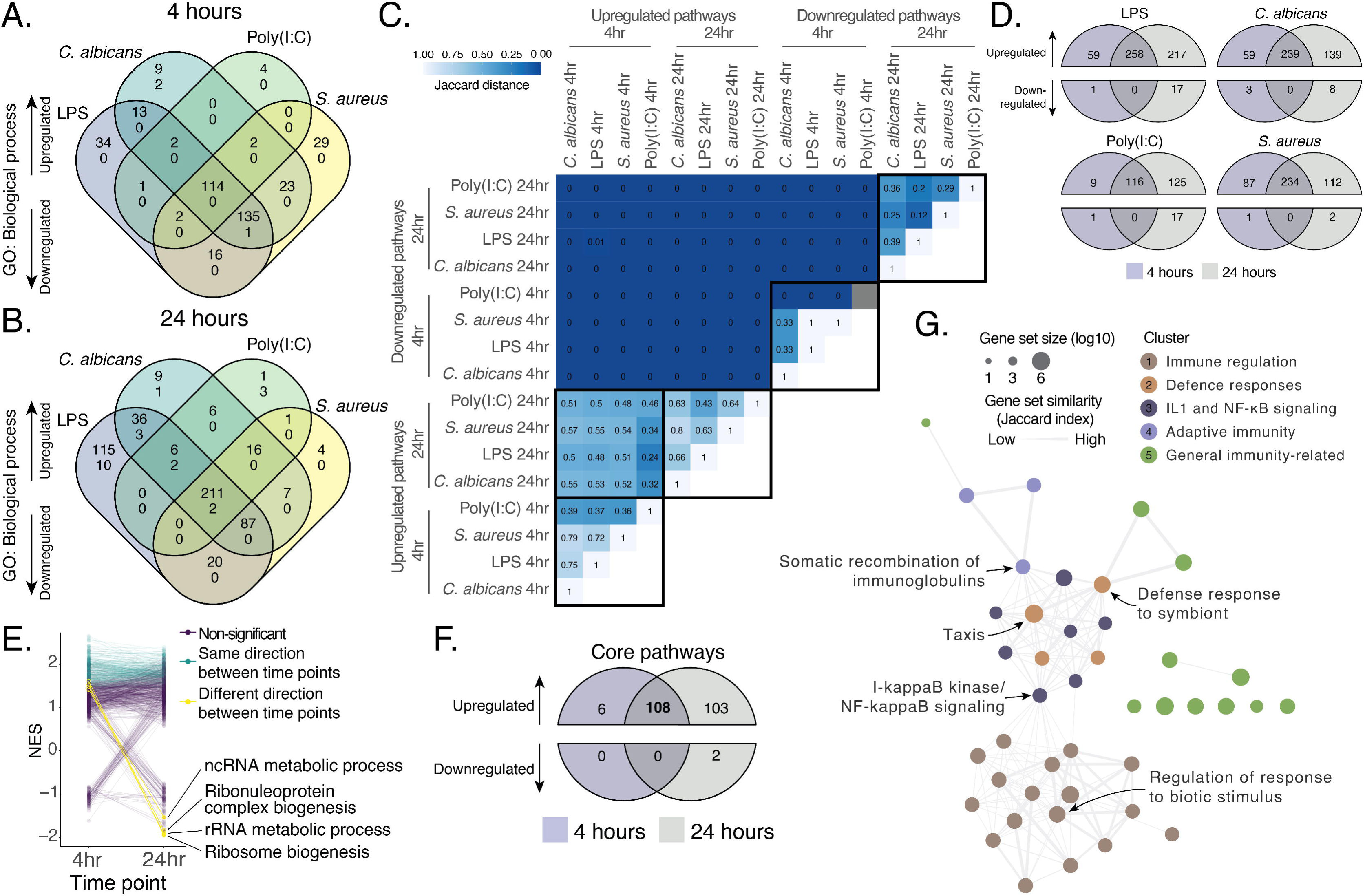
*Pathway analysis.* A-B) Overlapping up- and downregulated gene sets per stimulus condition, as analyzed by GSEA upon A) 4 hours stimulation and B) 24 hour stimulation. C) Heatmap of the Jaccard similarity index of the set of pathways up- or downregulated in each of the stimuli at each time point. D) Overlapping up- and downregulated pathways at both time points per stimulus condition. E) Comparison of pathway enrichment for 4- and 24 hour LPS stimulation, highlighting gene sets upregulated at 4 hours stimulation and downregulated at 24 hours stimulation, as indicated by the Normalized Enrichment Score (NES). F) Overlap of gene sets induced by all stimuli at both time points. G) Network diagram of core gene sets upregulated as a result of all four stimuli at both time points. The size of each point indicates the gene set size. The colors indicate the five clusters by similarity (Figure S4). Edge weight indicates the semantic similarity between gene sets.

Comparing the upregulated genes at both time points per stimulus revealed substantial overlap (Figure 2C-D, Table S26). Specifically for LPS, we find an overlap of pathways in RNA metabolism such as ncRNA and rRNA metabolism and ribosome- and ribonucleoprotein biogenesis, which are upregulated at 4 hour LPS stimulation and downregulated specifically after 24 hour LPS stimulation (Figure 2E and S12B-E, Table S26), potentially reflecting our previous findings on intron retention losses in splicing- and RNA processing-related genes (19). This highlights mechanisms of post-translational regulation in the immune response and indicating these mechanisms may be induced upon a wider variety of pathogen stimuli than described previously (74–76).

### Gene sets universally induced upon pathogen stimulation reveals a core network of immune stimulation and regulation

To identify core functions in the immune responses to stimuli derived from different pathogens, we overlapped the induced pathways for all four stimuli at both time points. We find 108 upregulated pathways shared by all stimuli at both time points (Figure 2F, Table S5 and S26), which we summarized giving representative subset of 42 gene sets (33). These form five clusters, representing functions in immune regulation, defense responses, IL1- and NF-κB signaling, adaptive immunity and other general immunity-related functions (Figure 2G and S13 Table S7). These pathways to form a two-sided network of immune-activating and immune-regulating pathways that is bridged by IκB kinase- and NFκB signaling (Figure 2G and S13, Table S7), highlighting its role in immune activation and regulation (79,80).

We find 2 downregulated gene sets specific to the core 24 hour response, with functions in transcription and translation (protein-DNA complex assembly and cytoplasmic translation), which partially aligns with the previously observed a downregulation of RNA metabolism- and translation-related gene sets upon LPS stimulation (Figure 2E, Table S26). These results indicate a shared downregulation of genes involved in translation upon pathogen stimulation, potentially reflecting expression regulation as a response to pathogens (Figure 2E-F, S11A-B and S12B-E, Table S26) (74,75).

### Genes induced by pathogen-stimulation are enriched for functions in immunity

We then opted to analyze our data on the gene level through a differential expression analysis. We analyze the differentially expressed genes (DEGs) upon 4- and 24 hour pathogen stimulation (Figure S14A, Table S8, Additional file 1), which we validate using publicly available RNAseq data of *in vitro* pathogen-stimulated PBMCs (39–42). This indicated a significant overlap in DEGs and correlations between gene-level log2 fold-changes identified from matching stimuli and time points (Figure S15, Table S9, Additional file 1). We have developed an interactive app for the exploration of our differential expression results available at emilvorsteveld.shinyapps.io/app_de.

Indicating the suitability of our approach to identify known genes involved in the innate immune response, we find genes involved in the toll-like receptor pathways to be upregulated at both time point, which includes toll-like receptor genes such as *TLR3* and *TLR4*, genes downstream the TLRs such as *IRF7* and *NFKBIA* and downstream signaling molecules such as *TNF*, *IL1B* and *IL6* (Figure S14D-E, Table S8). We find the genes in these pathways to have a higher differential expression at the 24 hour time point, while the highest outliers included signaling molecules downstream of these pathways (*IL6, IL1B, CXCL8* and *TNF*) were found upon the 4 hour time point (Figure S14F, Table S8).

Surprisingly, we find more DEGs downregulated upon 4- than 24 hour LPS stimulation (N=176 and N=149, Figure 3A). Early downregulated DEGs for LPS were enriched for functions in TNF production, which was distinct from the downregulated DEGs at the 24 hour time point. Examples include *CD84*, a suppressor of B- and T cell responses (81), *NLRP12*, a regulator of inflammation (82) and *NLRC4*, an inflammasome component that responds to bacterial components independent of the LPS-TLR4 pathway (83), indicating the early-onset downregulation of TNF-regulatory mechanisms upon LPS stimulation (Figure S14C, Table S27).

**Figure 3:**
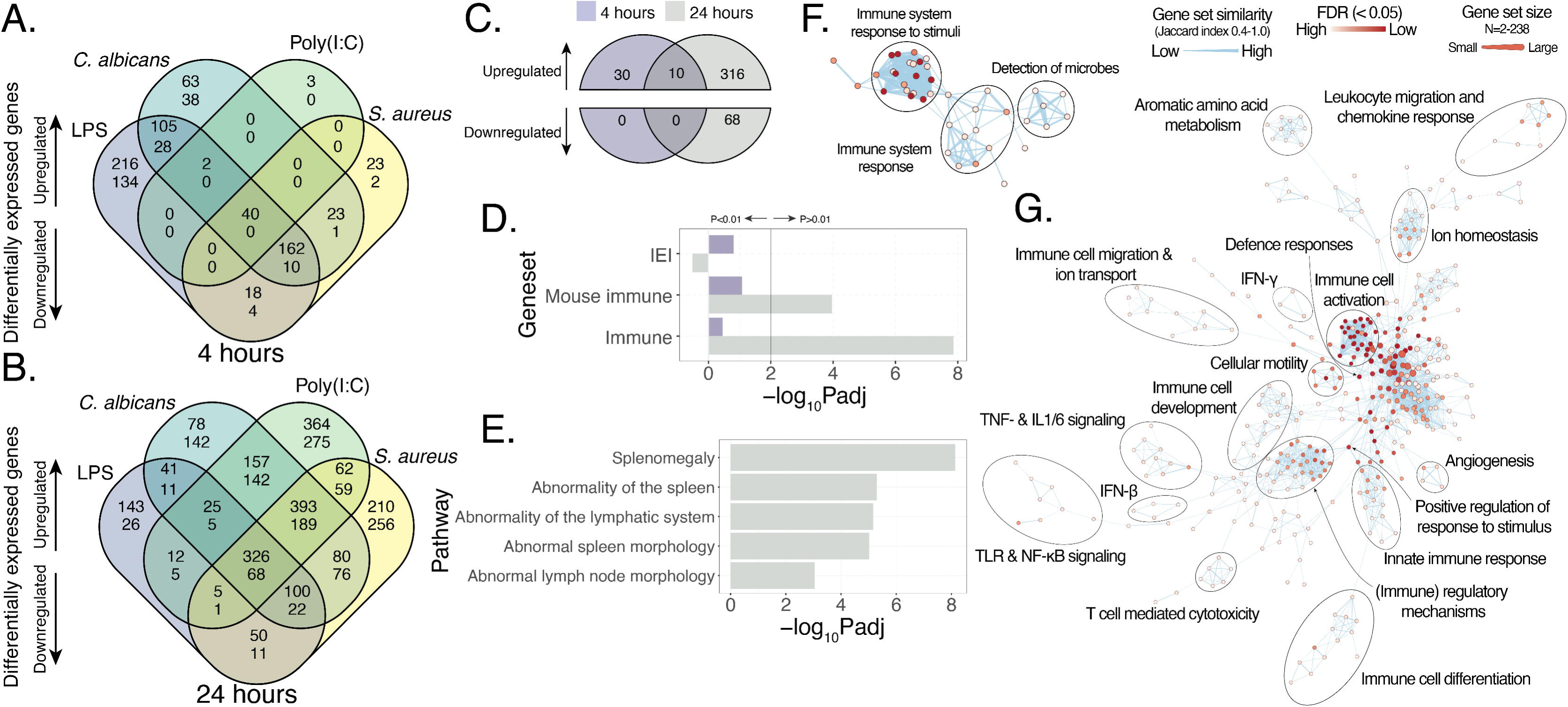
*Differential gene expression analysis.* A-B) Overlapping up- and downregulated DEGs at A) 4 hours stimulation and B) 24 hours stimulation. C) Overlapping up- and downregulated genes induced by all stimuli at either time point. D-E) Enrichment analysis of core differentially expressed genes at either time point using D) IEI genes and immunity-related genes in mouse (Mouse immune) and human (Immune) and E) using genes in the Human Phenotype Ontology, representing various human phenotypes. F-G) Enrichment map generated from protein-protein interactions of the genes upregulated as a result of all stimuli at F) 4 hours and G) 24 hours stimulation. Edges indicate similarity between gene sets (Jaccard index), node color indicates the enrichment P value and node size indicates gene sets size.

### Core genes induced by pathogen stimulation

We overlapped the upregulated DEGs identified in each of the stimulus conditions at both time points, revealing 40 shared DEGs at the 4-hour time point and 394 shared DEGs at the 24-hour time point (Figure 3A-B, Table S8 and S28, Additional file 1). We find 30 core DEGs specific to 4-hour pathogen stimulation (4.59%; Figure 3C, Table S28), with functions in the immune response to viruses (e.g. *OAS1-3* and *ISG15*, Table S8 and S28) and cytokine- and interferon responses (e.g. *IL1RN* and *USP18;* Table S8 and S28). Core DEGs specific to the 24-hour time point (N=316) were enriched for functions in NF-κB signaling (e.g. *NFKBIA* and *RIPK2*), and TLR signaling (e.g. *HCK*) and cytokine responses (e.g. *STAT1*). We find IEI genes to core DEGs at both time points including *IL1RN* and *IFIH1* (51.1% overlap at 4 hours and 17.8% at 24 hour stimulation; Figure 3C, Table S8 and S28). We find 68 core genes downregulated upon 24 hour stimulation, with plasma membrane- and vesicle-related functions (Figure 3B-C, Table S8 and S28). Overlapping the upregulated DEGs shared by all stimuli at both time points yields 10 genes, which are enriched for genes with functions in viral defense responses such as *IFIT3* and *IFI44L* (84,85) and genes with interferon-regulatory factor (IRF) transcription factor motifs, reflecting the direct transcription of IRF genes upon TLR stimulation (86). However, these shared DEGs are mainly dependent on the viral stimulus, which resulted in the fewest DEGs (Figure 3A-B and S14A, Table S12 and S28).

### Core genes induced by pathogen stimulation are enriched for genes associated with immune phenotypes and immunodeficiency

Core DEGs at the 24 hour time point are enriched for genes with functions in immunity in both human and mouse (Figure 3D, Table S12) and for genes associated with phenotypic traits such as abnormalities of the spleen and lymphatic system such as splenomegaly and abnormal lymph node morphology, potentially reflecting adaptive immune-related signaling occurring upon pathogen stimulation (Figure 3E, Table S12). While we do not find an enrichment of genes known to be involved in IEI, we find 8 and 20 core upregulated DEGs to be associated with IEI, respectively (20% and 6.1%), which share *IL1RN* (Figure 3C-D, Table S13 and S28). Considering all DEGs (N=571), we find 29 genes known to be responsible for an IEI characterized by infectious disease, which constituted a significant enrichment (0.81%, Fisher’s exact P=0.0018, Figure 3C and S14G, Table S13).

### Pathogen stimulation DEGs are enriched for genes associated with complex and immune-mediated diseases and genes associated with immune-cell specific eQTLs

We performed an enrichment analysis of genes associated with publicly available GWAS trait-associated genes among the pathogen-induced DEGs (Figure S7B, Table S14) (52). This revealed substantial overlap, with 10 shared enriched gene sets for all stimuli at both time points (0.42%, Figure S7C, Table S14), including traits involved in immune- and blood cell counts and with complex- and immune-mediated diseases such as psoriasis, systemic lupus erythematosus and type 2 diabetes, potentially reflecting the overlapping genetic etiology of these complex immune disorders (Figure S7D, Table S14) (87). We further find an enrichment of genes associated with Alzheimer’s disease and neuroticism, likely reflecting immune-mediated mechanisms in these disorders and traits (88,89).

We then investigated the overlap of our pathogen-response DEGs with expression quantitative trait loci (eQTL)-associated genes (eGenes) in immune cell types (32). We find an enrichment of a substantial number of eGene sets for all stimuli, while for instance 4- hour Poly(I:C) stimulation only resulted in an enrichment of T cell-related eGenes. We find a particularly high enrichment of eGenes associated with M2 macrophages and monocytes (OR range 3.14-3.80 and 3.13-3.72; Figure S8B, Table S15). These results indicate substantial overlap of our pathogen-stimulation DEGs with a broad set of immune cell-associated eGenes, highlighting T cell and monocyte-specific effects in particular.

### Pathogen stimulation induces a transcriptional response of naive CD4+ T cells and monocytes

To infer the cell type-specific responses that occur upon pathogen stimulation, we investigated the differential expression of genes uniquely expressed in 15 immune cell types in our pathogen stimulation experiments (32). We find genes specifically expressed in naïve CD4+ T cells and classical monocytes to be upregulated upon the 4 hour *C. albicans*, LPS and *S. aureus* stimulation and for 24 hour *C. albicans* and *S. aureus* stimulation, indicating a response of these cells in the response of *in vitro* pathogen stimulation. We find an upregulation of non-classical monocyte-specific genes upon 24-hour pathogen stimulation, likely reflecting the lack of expression of *TLR3* in human monocytes, in contrast to *TLR2* and *TLR4* (90,91). Surprisingly, NK cell-specific genes were downregulated upon 4 hour *S. aureus* stimulation and for all stimuli after 24 hours of stimulation (Figure S16A, Table S6), which were enriched for functions in leukocyte-mediated cytotoxicity (Figure S16B). This may indicate the induction of regulatory mechanisms at this late stimulation time point, with for instance immunoregulatory cytokine *IL10* upregulated upon 24-hour pathogen stimulation (Table S8) (92).

### Network analysis indicates common functions of differentially expressed genes between pathogen conditions

To gain insight into common functions of DEGs upregulated after all 4 or 24 hour of microbe stimulations (N=40 and N=326, Table S10), we visualized the biological functions of these genes using an enrichment map (30). We find a small network for the core DEGs at the early time point, highlighting biological functions involved in immune system processes such as the recognition of microbial stimuli and immune response-related gene sets that are subsequently induced (Figure 3F, Table S16-17). In contrast, the core DEGs at the late time point form a much larger pathway network, representing a wider range of biological functions. These include immune system processes, such as IFN-γ responses, with immune cell activation-related gene sets forming a central highly enriched and interconnected core of this network. This network includes both immune system activation, differentiation and regulation but also metabolism-, homeostasis-, migration- and transport-related pathways. This result reflects the pathway analysis of the 24 hour stimulation data, which included a wide variety of biological processes that were found to be upregulated upon pathogen stimulation. We find highly enriched nodes such as defense responses and the positive regulation of response to stimuli in this core gene network to link multiple larger clusters (Figure 3G, Table S18-19). These networks indicate the core functions of gene expression induced upon *in vitro* pathogen stimulation, which induces a small set of early-response genes, with a broader set of genes induced over time.

### Genes without a known function are differentially expressed upon in vitro pathogen stimulation

Following the finding that our pathogen stimulations support the broad and specific expression of genes and pathways related to immunity and to the specific pathogen stimuli respectively, we analyze the expression of genes that have not previously been associated with any function in immunity in our dataset. To this end, we classified all DEGs by their functional associations; genes with roles in immunity, with genes associated with IEI separately. We further classified protein-coding genes without a well-known function (henceforth “uncharacterized genes” and “characterized genes”) using the “annotation score” in UniProt (>2, characterized; ≤2, uncharacterized). As a quality control measure, we assessed the number of publications associated with the characterized and uncharacterized DEGs we identified using our pathogen stimulations using a public database of understudied genes (Figure S6A-B, Table S11) (11). We find the characterized DEGs we identified to be associated with significantly more publications than the uncharacterized DEGs (mean 66.2 and 1.4; Wilcoxon rank sum test p < 2.2*10^-16^, Figure S6C, Table S11), supporting the notion that these genes are unlikely to be associated with immunity-related processes already.

Considering each condition-time point combination separately, we find on average 6.5% of DEGs to be associated with IEI (range 4.0-17.8%, 177 genes) and an average of 34.5% of genes having with a role in immunity (range 25.2-51.1%, 827 genes). We find an average of 1.3% of DEGs to be uncharacterized (range 0.0-2.0%), yielding 78 genes that were differentially expressed in at least one condition (Figure 4A and S14A, Table S10 and S28). Of these genes, 13 were differentially expressed at the 4 hour time point and 72 at the 24 hour time point. We find five uncharacterized genes to be differentially expressed in all conditions at the 24 hour time point (6.9%; *C11orf21*, *SMCO2*, *NRIP3*, *C1orf122* and *ITPRIPL*), with an additional gene being differentially expressed upon all stimuli without considering the time point, resulting in six genes (7.7%; includes *KIAA0040*) (Figure 4B, Table S10 and S28).

**Figure 4:**
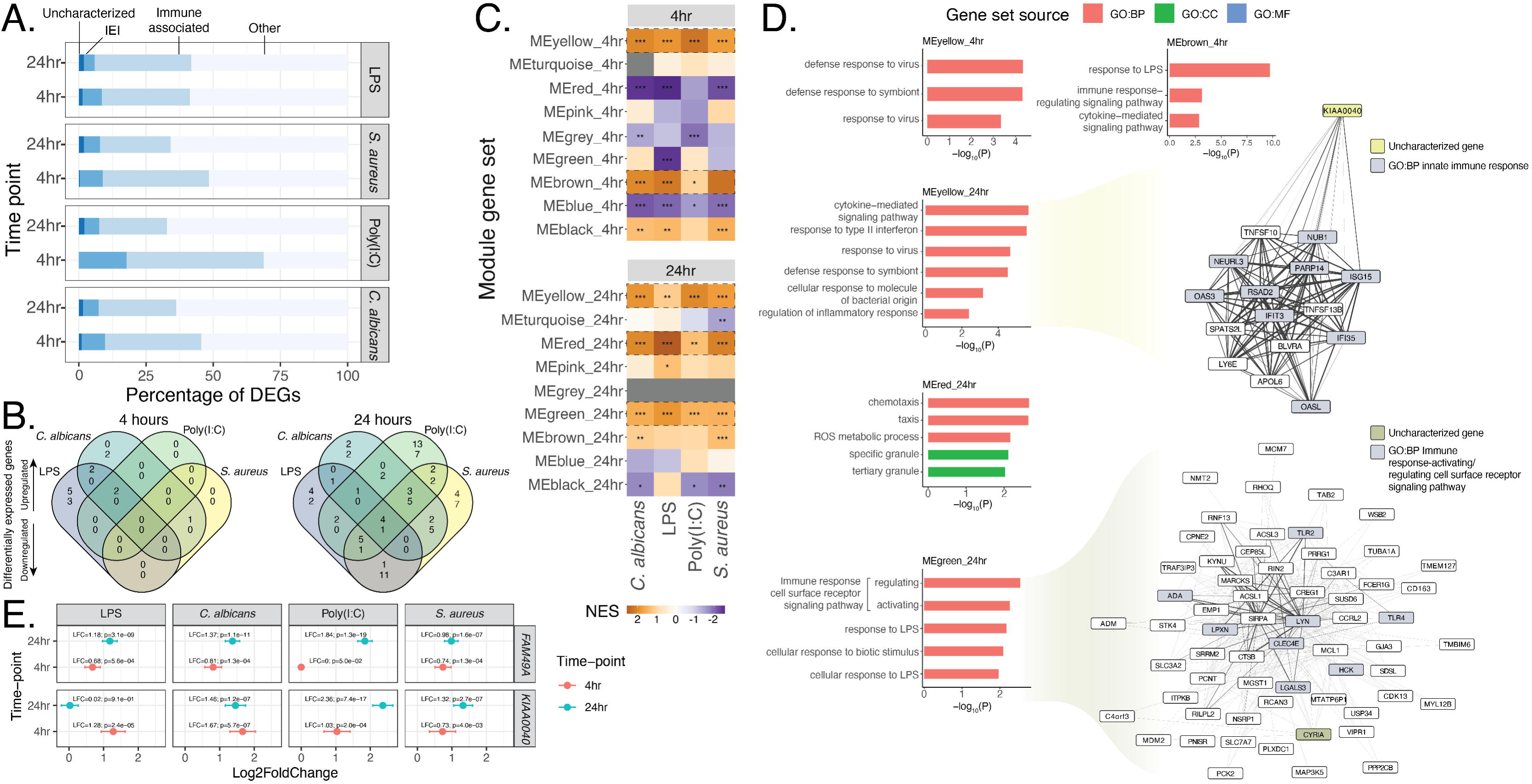
*Identification of uncharacterized genes differentially expressed upon microbial stimulation.* A) Functional classification of the DEGs identified per time point and stimulus condition, identifying genes with known (immune) functions and those without a previously described function. B) Overlap of uncharacterized DEGs identified for each stimulus condition at either time point. C) Gene set enrichment analysis using the gene modules identified through the co-expression analysis, indicating the normalized enrichment score and the adjusted p value. Modules that were further investigated using a functional enrichment analysis are highlighted with dashed boxes. E) Functional enrichment analysis of the co-expression modules that include uncharacterized genes that were found to be upregulated upon all stimuli. For the 24 hour yellow and green modules, the highest correlating genes of the uncharacterized gene in each module are highlighted in yellow, together with genes in the network with functions related to the innate immune response and immune cell signaling in purple. E) Differential expression of uncharacterized genes *KIAA0040* and *CYRIA (FAM49A)*. Asterisks indicate level of statistical significance: * p ≤ 0.05, ** p ≤ 0.01, ***p ≤ 0.001. LFC, log2 fold-change; NES, Normalized enrichment score.

### Co-expression analysis identifies gene modules associated with functions in the immune response

We were motivated by the uncharacterized genes differentially expressed upon pathogen stimulation and sought to identify potential biological functions to these genes using a co-expression analysis, which allows for the identification of guilt-by-association predictions of gene function (93). We used the stimulated and control data for each time point separately to generate weighted gene co-expression network (Figure S9A-G). We identified 9 co-expression gene modules at either time point (mean 537.3 genes at 4 hours, range 83-2073; 575.9 at 24 hours, range 47-2278, Figure S17A, Table S20). We performed a pathway analysis of the DEGs found through our pathogen stimulation experiments using each of the gene modules at the matching time point to validate that the genes in each co-expression module are induced upon *in vitro* pathogen stimulation. This yielded six modules that were differentially expressed in all stimulus conditions, with three at either time point (Figure 4C, Table S21). At the 4 hour time point, the yellow and brown modules were upregulated for all stimuli (Figure 4C). This module is enriched for genes with functions in viral responses and the response to LPS and cytokine pathways, respectively (Figure S17B, Table S22). At the 24 hour time point, we find the yellow, red and green modules to be differentially expressed in all stimulus conditions (Figure 4C). We for instance find the red module at the 24 hour time point to be enriched for genes with functions in chemotaxis, ROS metabolism, granule-related functions and cytokine receptor binding (Figure S17C, Table S22). We find 39 uncharacterized DEGs to be part of these co-expression modules. Here, we highlight *KIAA0040* and *FAM49A*, which are part of the yellow and green modules generated using the 24 hour-stimulated samples, respectively (Figure 4D).

### A co-expression module that includes uncharacterized gene KIAA0040 involved in innate immune responses

The yellow co-expression gene module found at the 24 hour time point to be enriched for immune-related genes involved in cytokine- and interferon type II signaling, responses to pathogens, including bacteria and viruses and the regulation of inflammatory responses such as *ISG15, OASL, OAS3* and *IFIT3* (Figure 4D and S17C, Table S20-21). This gene module is upregulated in all stimulus conditions includes uncharacterized gene *KIAA0040* (Figure 4C-D, Table S8 and S20). The expression of *KIAA0040* is most highly correlated with genes with functions in the innate immune response such as *IFIT3* and *OAS3* (Figure 4D)*. KIAA0040* was recently found to promote glioma formation through the JAK2/STAT3 pathway (94) and is a member of a gene expression module that responds to IL7 stimulation in primary Sjögren syndrome patients (95). Investigating *KIAA0040* in single cell expression data of blood cells indicates its expression in erythrocytes, B cell lineages and neutrophils (Figure S18A, Table S23) (57,58). Similarly, gene expression in sorted PBMCs indicates that *KIAA0040* is expressed in neutrophils, memory- and naïve B cell lineages (Figure S18B-D, Table S24) (32,55,59,60).

### A co-expression module that includes uncharacterized gene CYRIA (FAM49A) involved in TLR signaling

The red co-expression gene module at 24 hours is enriched for genes involved in chemotaxis, reactive oxygen species production, granule formation and the regulation and activation of immune responses through cell surface receptor signaling and responses to bacterial stimuli such as LPS, which includes genes such as *TLR2, TLR4, ADA,* and *LYN* (Figure 4D and S17C, Table S20-21). This gene is induced by multiple stimuli at the 24 hour time point and includes uncharacterized gene *CYRIA* (also known as *FAM49A;* Figure 4B and E, Table S8 and S20). We find the core of this network to includes various genes involved in TLR2- and TLR4 signaling, such as downstream signaling molecules *LYN* and *ADA* (Figure 4D, Table S20).

*CYRIA* codes for CYRI-A, which is part of the CYRI protein family that share functions in the regulation of cellular motility and migration through protrusion generation (96). It shares 80% sequence homology with its paralogue *CYRIB,* which regulates cytoskeletal remodeling in phagocytosis, thereby protecting against bacterial infections (97). CYRI-A is a key regulator of micropinocytosis dependent on PI3K and RAC1 (98). An investigation of single cell expression data of blood cells reveals high expression of *CYRIA* in CD14^+^- and non-classical monocytes (18.1% and 18.4% expressed) and in progenitor cells and B cells (Figure S18A, Table S23) (57,58). Evaluation of the expression of *CYRIA* in sorted PBMCs indicates its expression in various cell types of the innate and adaptive immune system including neutrophils, monocytes, NK cells and B cells (Figure S18B-D, Table S24) (32,55,59,60).

### The expression of uncharacterized genes is correlated with immune genes in publicly available gene expression data

We sought to validate the results from our co-expression analysis using publicly available co-expression data. We performed a pathway enrichment analysis using the gene-gene correlation coefficients for the uncharacterized DEGs we identified (61,62). We find the genes co-expressed with *KIAA0040* to be enriched for functions in adaptive immune responses such as B cell signaling and immunoglobulin recombination, which is concordant with the expression pattern of *KIAA0040* (Figure S18, Table S23-25). We further find an enrichment of genes involved in cell surface receptor signaling and cytokine signaling (Figure S19A, Table S25), further illustrating the candidate function of *KIAA0040* in cytokine signaling in innate and adaptive immunity. Genes co-expressed with *CYRIA* are enriched for functions in immune cell signaling, leukocyte function and migration, T cell responses and phagocytosis (Figure S19B, Table S25). While this result is not completely concordant with the expression pattern of *CYRIA,* which is highly expressed in monocytes (Figure S18A-D, Table S23-24), these results provide support for *CYRIA* as candidate regulator of phagocytosis and chemotaxis during immune responses to bacteria, which is initiated by stimulation of TLR2 and TLR4, indicating a candidate function in both innate and adaptive immune cells.

### Uncharacterized genes differentially expressed upon pathogen stimulation provide candidate variants in patients with inborn errors of immunity

A majority of patients with rare immune-mediated disease such as IEI remain without a molecular diagnosis. We therefore postulate that there remain genes currently undescribed as causative for these disorders. We analyzed rare variants in the uncharacterized genes we identified to be differentially expressed upon our pathogen stimulations in a cohort of >1,300 IEI patients that underwent diagnostic whole exome sequencing (63). We identified 901 variants and prioritized 5 variants in 4 uncharacterized genes that were rare or absent in population databases in previously undiagnosed IEI patients with recurrent infections (Table S29-30) (65,66). Of these, we find 4 missense variants with a high predicted impact on the protein (CADD PHRED range 23-27.3), which were also phylogenetically conserved (PhyloP range 2.0-9.8). We identified missense variants in *YPEL1* in two patients with chronic (fungal) infections. This gene is differentially expressed in response to multiple stimuli and is constrained against loss-of-function variation (gnomAD LOEUF 0.77, Table S8 and S30) and is part of a co-expression module enriched for genes associated with the response to LPS, including cytokine responses and immune regulation (Figure 4D, Table S20-21). We also identified a frameshift variant in *TMEM185B* in a patient with suspected dyskeratosis congenita with recurrent infections and stomatitis and sparse hair. *TMEM185B* is differentially expressed in response to LPS and is constrained against loss-of-function variation (gnomAD LOEUF 0.77, Table S8 and S30) and was part of the same co-expression module as *KIAA0040* (MEyellow 24hr, Figure 4C-D, Table S20). These variants provide potential causes of IEIs associated with a susceptibility to infectious diseases.

## Discussion

The identification of new disease genes associated with rare genetic disorders is vital for the diagnosis and functional characterization of these diseases. This is hampered in part due to the incomplete functional annotation of genes, complicating variant interpretation (2).

Studying immune-mediated diseases using transcriptomics is a very useful approach, as the affected tissue is readily available and the cells can be stimulated *in vitro,* providing a method for mimicking *in vivo* infections. We applied RNA sequencing to profile the transcriptome of *in vitro* pathogen stimulated primary immune cells (PBMCs) for the identification of novel candidate genes in innate immune responses. We analyzed the transcriptional response on gene- and pathway level and explore the potential functions of previously uncharacterized genes in innate immune responses using co-expression analysis.

We describe a set of pathways that are found to be upregulated as a result of all pathogen stimuli, highlighting a network of immune-stimulating and -regulating processes. We highlight rRNA and ncRNA metabolism as mechanisms that are upregulated at early LPS stimulation but downregulated with a longer stimulation time. We describe core pathways that are upregulated in response to all stimuli to be characterized by immune-related pathways with functions in both innate and adaptive immunity. We found the number of DEGs in each stimulus condition to be highly variable, where for LPS stimulation a much larger number of DEGs at the early time point was found. While this stimulus is the most widely used among publicly available RNAseq datasets of human tissue samples (511, 110, 40 and 62 datasets of human samples for LPS, Poly(I:C), *C. albicans* and *S. aureus,* respectively; retrieved in the Gene Expression Omnibus on February 17th 2025 (38)), our results may indicate this stimulus may not be representative for other pathogens. An analysis of cell type-specific immune genes identified a specific response of T cell- and monocyte-related genes, indicating a response of these cell types to our pathogen stimuli. We further found a downregulation of cytotoxicity genes associated with NK cells, potentially indicating regulatory mechanisms. An overlap with eQTLs identified in various immune cell types further highlights the response of these cell types together with a broad induction of immune cell expression. We find the DEGs to be enriched for genes associated with GWAS hits related to immune mediated diseases and phenotypes. Together these highlight overlap with immune cell expression and complex traits and disorders in gene expression induced upon the pathogen response. The suitability of our approach for the use of gene expression as a proxy for protein prevalence can be seen from the expression of genes encoding for cytokines such as *TNF*, *IL1B* and *IL6*, which are commonly studied by more classical protein assays (e.g. ELISA), and are well established for these response (99). Our data further indicate an LPS-specific downregulation of regulatory factors of this pathway. The differential gene expression and pathway enrichment results from the current study can be interactively explored and downloaded at emilvorsteveld.shinyapps.io/app_de.

We have identified 78 genes without a previously described function to be differentially expressed in response to our pathogen stimuli. We found 26 uncharacterized genes (33.3%) to be co-expressed with sets of other genes, potentially indicating the biological network these genes play a role in. We highlight *KIAA0040* and *CYRIA* (*FAM49A*), which we found to be co-expressed with genes that play a role in the innate immune response and in downstream signaling from the PRRs (TLR2 and TLR4), respectively. We analyze rare genetic variants in these uncharacterized genes in a large cohort of IEI patients (63), identifying 5 candidate variants that are prioritized for further follow-up. These variants may lead to the identification of novel genes underlying IEI and provides an example where transcriptomics aids in transcriptome-based gene discovery in rare disease.

There remain various obstacles to overcome for further characterization of genes without a known function. Transcriptomics plays an important role in these endeavors (100). Long-read sequencing can be used for the accurate identification of transcripts expressed during an immune response, including transcripts that are difficult to detect using conventional RNAseq methods, providing a more accurate view of the transcriptome and leading to further understanding of the human immune response (19,101,102). Single cell sequencing provides data on the specific cells or cell types that express a gene or transcript (103). This has led to further characterization of the immune response at a previously unprecedented level of detail. Recent advances in methodology allow for the combined application of long read technologies using single-cell sequencing libraries, providing full-length transcript information at the single cell level (104). Further efforts using these methods in the field of human immunology will allow for the identification of additional genes and transcripts with functions in biological mechanisms including immune-related pathways.

The identification of new disease genes is a major driver for additional diagnostic yield for patients with rare genetic disorders (2,63). Here, we describe the identification of 5 candidate variants in 4 genes without a previous function in patients with inborn errors of immunity. These variants will be further followed up on to assess their role in these rare diseases. Transcriptomics has played an important role in the identification of phenotype-genotype associations in health and disease, leading to the identification and validation of candidate disease genes. Transcriptomics has been shown to be effective in a clinical setting (6). Immune-mediated diseases such as IEIs provide a unique opportunity for transcriptomics, as the affected tissue is easily accessible and because an infection can be mimicked *in vitro*. Further studies using PBMC stimulations in cohorts of or even individual patients with rare immune diseases, comparing the transcriptional response with a healthy population could therefore be promising to provide additional diagnostic yield through the investigation of differential splicing and expression patterns. Our identification of rare variants in uncharacterized genes could further provide additional diagnostic yield and could expand the currently known disease genes in this field.

### Limitations of the study

This study has several limitations. First, the extent to which conclusions can be drawn concerning the human immune response is limited by our sample size. While we have shown that most of the variability in the transcriptional response is related to the stimulation time and the condition, this explorative study is mainly meant to provide novel mechanisms and candidate genes that may be suited for functional follow-up and prioritization in the investigation of genetic variation in rare immune-mediated disease. There exist inter-individual differences in the human immune response, dependent on age, sex and genetic differences. This study was done in a population of European ancestry, potentially limiting the extrapolation of our findings to other populations. Our small-scale study was underpowered to further investigate these interindividual differences in the immune response (73). Future studies could investigate these influences on the immune response using a larger sample size. Second, while PBMCs represent a mixed population of peripheral immune cells, allowing for communication across cell types after *ex vivo* pathogen stimulation, these experiments may not fully represent an *in vivo* immune response (105). Third, in this study we have generated bulk transcriptomics data, which does not provide transcriptional information on the level of cell types. Future studies could employ single-cell sequencing of pathogen-stimulated immune cells to more completely ascertain the cell types that express the genes and gene modules we identify here, which has for instance implicated *CLEC12A* as an IFN-regulated gene (106). Additionally, novel sequencing methods such as long read sequencing have been employed to study transcript expression in the human immune response, which could be adapted to allow for single-cell transcript expression analysis (19,103,104). Fourth, our analysis of rare variants in uncharacterized genes is limited by the sample size of the patient cohort and by technical limitations in the sequencing, as the used exome capture kits have limited probes targeting certain genes.

Finally, as is a general limitation of explorative omics studies, further functional follow-up of the candidate associations we identified is necessary to confirm these functional associations to characterize these genes.

## Declarations

### Ethics approval and consent to participate

PBMCs were retrieved form healthy, anonymized donors, as part of the human functional genomics project (HFGP). The HFGP study was approved by the Ethical Committee of Radboud University Nijmegen, the Netherlands (no. 42561.091.12). Experiments were conducted according to the principles expressed in the Declaration of Helsinki. Samples of venous blood were drawn after informed consent was obtained.

### Consent for publication

Not applicable.

### Availability of data and material

Raw sequencing data is available on EGA under accession number EGAS50000000007. Scripts used to generate the results described in this paper can be found at github.com/EmilVorsteveld/quantseq_pbmc.

### Authors’ contributions

Conceptualization: EEV, SK, MGN, AH; data curation: EEV, SK, CK; formal analysis: EEV, SK, CK; investigation: EEV, SK; resources: ; software: CK; supervision: PtH, MGN, AH; visualization: EEV; writing—original draft: EEV; writing—review and editing: All authors. All authors read and approved the final manuscript.

## Acknowledgement

We would like to acknowledge colleagues from the Radboud Genomics Technology Center for their support with sequencing and colleagues from the Department of Internal Medicine for their support with our PBMC stimulation experiments.

## Supplemental material

- Figures S1-S19: Supplemental figures
- Tables S1-S29: Supplemental tables
- Supplemental file 1

## Supplemental figures

Figure S1: *Gene body coverage.* Sequencing coverage across the gene body per positional percentile of 3802 housekeeping genes for the included samples.

Figure S2: *Expression correlation matrices*. Comparing the correlation of gene expression levels in each of the control (RPMI) sequencing libraries. Each individual is indicated with a letter, where “repl” indicates a replicate sequencing libraries (technical replicates) and the numbers “1” and “3” indicate separate control samples (biological replicates). A) Correlation between technical replicates of 4 hour control samples. B) Correlation between technical replicates of 24 hour control samples. C) Correlation between biological replicates of 4 hour control samples. D) Correlation between biological replicates of 24 hour control samples.

Figure S3: *Elbow plots for k-means clustering.* A) Elbow plot to select the number of gene clusters. B) Elbow plot to select the number of sample clusters.

Figure S4: *Elbow plots for k-means clustering of the summarized core pathways.* Selection of the optimal number of clusters of the core upregulated pathways.

Figure S5: *Volcano plots.* Volcano plot per condition and time point combination. We defined a gene DE when the adjusted P value was <0.01, with upregulated DEGs having a log2 fold-change ≥ 1 and downregulated genes ≤-1. Upregulated genes are indicated in green, downregulated genes in purple.

Figure S6: *Number of publications associated with characterized and uncharacterized genes.* A-B) Overlap of the genes available in the Find My Uncharacterized Genes (FMUG) database with A) All genes and DEGs in our dataset and B) separately for characterized and uncharacterized DEGs. C) Violin plot showing the number of publications associated with characterized and uncharacterized DEGs as registered in FMUG (11).

Figure S7: *GWAS gene set enrichment.* A) The number of genes associated with GWAS traits., binned by N=0, N=1, 1>N≤10, 10<N≤100 and 100<N≤1,000 and N<1000 genes. B) The number of enriched gene sets per condition-time point combination. C) An upset plot showing the overlap of enriched gene sets associated with GWAS traits per stimulus and time point. D) Odds ratios for 10 GWAS gene sets that were enriched for all stimulus-time point combinations.

Figure S8: *Enrichment of genes associated with immune cell eQTLs.* A) Upset plot showing the overlap between gene sets mapped to cell type-level eQTLs. The top 20 largest overlaps are shown. B) Odds ratios calculated for the overlap of the pathogen stimulation DEGs with genes harboring eQTLs at stimulus and time point. Asterisks indicate level of statistical significance: * p ≤ 0.05, ** p ≤ 0.01, ***p ≤ 0.001.

Figure S9: *Weighted gene co-expression network analysis.* A-C) Sample trees constructed from expression data from A) both time points B) 4 hour-stimulated samples C) 24 hour-stimulated samples. D-G) Soft cutoff choice and mean connectivity for WGCNA using a Scale free topology model fit for D-E) 4 hour-stimulated samples and F-G) 24 hour-stimulated samples.

Figure S10: *Principal component analysis per stimulation time point.* Comparing gene expression between the samples per time point using principal component analysis using A) 4 hour stimulated samples and B) 24 hour stimulated samples. Shapes indicate the donors and colors indicate the condition.

Figure S11: *Gene set co-regulation analysis.* Gene Ontology Biological Process gene sets that are highly correlated across samples. Generated using a gene set co-regulation analysis (GESECA). A) Gene sets ordered by adjusted P value. The scaled expression of the genes in each gene set is plotted per sample. B) Expression patterns for a selection of gene sets across samples.

Figure S12: *Gene set enrichment analysis.* A) Number of up- and downregulated gene sets per stimulus and time point. B-E) GSEA plots of gene sets upregulated at 4 hours and downregulated after 24 hours of LPS stimulation (Figure 2E): B) Ribosome biogenesis, C) ncRNA metabolic process, D) rRNA metabolic process and E) Ribonucleoprotein complex biogenesis.

Figure S13: *Clustering of summarized core pathways.* A) GO term similarity matrix (SimRel) from a representative set of 42 pathways generated using REVIGO from 108 pathways induced upon all pathogen stimuli at both time points, which form five clusters (Figure S4).

Figure S14: *Differentially expressed genes.* A) The number of differentially expressed genes (DEGs) identified per stimulus and time point colored by the functional classification of the genes. B-C) Functional enrichment of the DEGs B) upregulated upon 4 hour Poly(I:C) stimulation and C) downregulated upon 4- or 24 hour LPS stimulation. D-E) Visualization of the toll-like receptor pathway genes with associated log2 fold-changes per condition for D) 4 hour stimulation and E) 24 hour stimulation. F) Distribution of log2 fold-change values for toll-like receptor genes. The log2 fold-change values are binned indicating LFC≤0, 0<LFC≤1, 1<LFC≤2, 2<LFC≤5 and LFC>5. G) Core DEG enrichment of IEI genes associated with innate immune defects, as defined by the IUIS in IEI gene table 6 (1).

Figure S15: *Expression validation using publicly available data.* Principal component analysis and DEG overlaps comparing genes expression generated in this work together with publicly available data from four publications of RNA sequencing data from pathogen-stimulated PBMCs. Comparison with A-B) LPS-stimulated PBMCs (39), B-C) Poly(I:C)-stimulated PBMCs (40), *C. albicans*- and *C. auris-*stimulated PBMCs (41) and PBMCs stimulated with various fungal stimuli, including *C. albicans* (42).

Figure S16: *Enrichment analysis using cell type-specific genes.* A) The enrichment is displayed for all stimuli at either time point. B) Functional enrichment of downregulated NK cell genes. Asterisks indicate level of statistical significance: * p ≤ 0.05, ** p ≤ 0.01, ***p ≤ 0.001. NES, Normalized enrichment score.

Figure S17: *WGCNA modules.* A) Number of genes per co-expression module generated from the 4- and 24-hour stimulated samples. Functional enrichment of the gene modules identified using the B) 4 hour stimulated and C) the 24 hour stimulated data, displaying the gene sets from the Gene Ontology, Reactome, CORUM, the Human Protein Atlas and KEGG.

Figure S18: *Expression of KIAA0040 and FAM49A publicly available (single cell) RNAseq data.* We investigated the expression of *KIAA0040* and *FAM49A* across cell types in A-B) the CELLxGENE database and C-E) in publicly available datasets of gene expression data from cell-sorted immune cell populations (32,55,57,60).

Figure S19: *Gene set enrichment analysis of publicly available co-expression data.* GSEA analysis of the Pearson correlation coefficients of A) *KIAA0040* and B) *FAM49A (CYRIA)* using publicly available expression data in ARCHS4 (61,62). NES, Normalized enrichment score; pval, P value; padj, Adjusted P value.

## Supplemental tables

Table S1: Transcript counts of 4 hour-stimulated PBMCs.

Table S2: Transcript counts of 24 hour-stimulated PBMCs.

Table S3: Heatmap expression values.

Table S4: Gene set co-regulation analysis.

Table S5: Pathways enriched per time point and stimulus.

Table S6: GSEA of immune cell type-specific genes from DICE.

Table S7: Core pathways summarized with ReviGO.

Table S8: Differentially expressed genes per time point and stimulus.

Table S9: Validation of DEGs using publicly available RNAseq data from pathogen-stimulated PBMCs.

Table S10: Functional classification of DEGs per time point and stimulus.

Table S11: Number of publications per gene as registered in Find My Understudied Genes (FMUG).

Table S12: Functional enrichment of core genes upregulated at either time point.

Table S13: Enrichment of immune genes among core genes upregulated at either time point.

Table S14: Enrichment of gene sets harboring GWAS hits.

Table S15: Enrichment of gene sets harboring cell type-specific eQTLs.

Table S16: Nodes of the pathway network analysis generated from core DEGs at 4 hours.

Table S17: Edges of the pathway network analysis generated from core DEGs at 4 hours.

Table S18: Nodes of the pathway network analysis generated from core DEGs at 24 hours.

Table S19: Edges of the pathway network analysis at generated from core DEGs at 24 hours.

Table S20: Gene modules identified using WGCNA at either time point.

Table S21: Enrichment analysis with GSEA using WGCNA gene modules with matched time points.

Table S22: Functional enrichment of WGCNA co-expression modules using g:Profiler.

Table S23: Single cell expression of uncharacterized genes *KIAA0040* and *FAM49A (CYRIA)* in blood cells using publicly available data.

Table S24: Expression of uncharacterized genes *KIAA0040* and *FAM49A (CYRIA)* in sorted immune cells in publicly available expression data.

Table S25: Pathway enrichment using gene co-expression data for uncharacterized genes *KIAA0040* and *FAM49A* (*CYRIA*).

Table S26: Overlapping and distinct pathways by stimulus and time point.

Table S27: Functional enrichment of genes downregulated upon 4- or 24 hour LPS stimulation.

Table S28: Overlapping and distinct DEGs by stimulation and time point.

Table S29: Rare variant filtering strategy.

Table S30: Rare variants in uncharacterized genes identified in a cohort of patients with inborn errors of immunity.

