## Additional file 1 for "Transcription-based identification of uncharacterized genes in the human immune response"

##### Supplemental results

###### Differential expression validation

We compared our LPS-stimulated samples to a publicly available dataset of LPS-stimulated PBMCs (1). Using PCA, we observed a separation between control (RPMI) and LPS-stimulated samples and a cluster of LPS-stimulated samples across datasets (Figure S6A). We found a significant overlap between the DEGs (overlap N=268, Figure S6B; Fisher’s exact OR=7.03, P<2.2*10^-16^), with a high correlation in expression (R2=0.66).

We compared our Poly(I:C)-stimulated samples at the 24 hour time point with a publicly available dataset of Poly(I:C)-stimulated PBMCs (2). We observed a clustering of the Poly(I:C)-stimulated samples across datasets. In contrast, the control (RPMI) samples form two distinct clusters, indicating differences between the control samples of either dataset (Figure S6C). We found a significant overlap between the DEGs (overlap N=1028, Figure S6D; Fisher’s exact OR=5.61, P<2.2*10^-16^), with a high correlation in expression (R2=0.75).

We then compared our *C. albicans* samples to two publicly available datasets (3,4). We find the samples of the first dataset to only cluster based on stimulation time and not by condition (Figure S6E). We find the expression of the DEGs obtained from 4 hour stimulation with live *C. albicans* and *C. albicans* beta glucan to be highly correlated (R2=0.80 and R2=0.76), yielding a significant enrichment (overlap N=57 and N=75, Figure S6F; Fisher’s exact OR=28.4, P<2.2*10^-16^; OR=14.6, P<2.2*10^-16^), we could not determine a correlation with the DEGs obtained from mannan stimulation at this time point due to a lack of overlap (N=2, 66.7%, Figure S6F). The DEGs from 24 hour stimulation with live *C. albicans*, *C. albicans* mannans or beta glucans were weakly correlated (R2=0.078; R2=0.34; R2=0.047), but still yielded a significant enrichment (overlap N=416, N=30, N=157, Figure S6F; Fisher’s exact OR=3.89, P<2.2*10^-16^; OR=6.09, P=2.76e-14; OR=5.04, P<2.2*10^-16^).

We find the *C. albicans* samples of the second dataset form clusters based on the stimulation time point (Figure S6G). An additional dataset of *C. albicans*-stimulated PBMCs yielded a high correlation in expression (4hr R2=0.82; 24hr R2=0.76), yielding a significant enrichment in the DEG overlap at 4 hours (overlap N=283, Figure S6H Fisher’s exact OR=50.71, P<2.2*10^-16^) and 24 hours (overlap N=753, Figure S6H; Fisher’s exact OR=13.16, P<2.2*10^-16^). These results indicate that the expression data obtained from our PBMC stimulation experiments are generally in line with those from previously published data (Table S9).

###### Pathway-level analysis of the transcriptomic response to in vitro microbial stimulation

Pathway analysis revealed 477 shared enriched gene sets between conditions and 124 unique gene sets (Figure 2A-B, Table S5). We generally find a higher number of enriched pathways upon 24 hour pathogen stimulation than at 4 hour stimulation (mean 368.5 and 266.5). At the 4 hour time point, we find the highest number of enriched gene sets for LPS stimulation (N=492) and the lowest for Poly(I:C) (N=125). We find an average of 1.67 and 8.5 downregulated gene sets at 4 and 24 hours, with up to 17 downregulated gene sets for 24 hour LPS stimulation (Figure 2A-B and S11A, Table S5).

Common and distinct pathways induced by in vitro microbial stimulation

Pathogen stimulation for 4 hours with our four stimuli resulted in a substantial overlap in upregulated pathways between the *C. albicans-*, *S. aureus-* and LPS stimulation (N=135, Table S5 and S26), with functions in immune-related functions such as IFN-γ production, cellular proliferation and the development and differentiation of immune cells. We find the pathways upregulated as a result of *C. albicans-* and *S. aureus* stimulation to be the most similar, followed by *C. albicans* and LPS. We find LPS stimulation to induce the highest number of distinct upregulated pathways (N=34), representing functions in translation, responses to reactive oxygen species and various other functions in cellular responses and development (Figure 2A, Table S5 and S26).

Comparing the pathways induced by all four stimuli at both time points, we find the highest similarity between time points for *S. aureus* stimulation (Jaccard index 0.54), whereas Poly(I:C) stimulation shows the lowest similarity (Jaccard index 0.46) (Figure 2C). We find the Poly(I:C)-induced pathways at both time points to be the most distinct from those in other stimulus conditions at the same time point (Jaccard index range 0.32-0.46 at 4 hours and 0.43-0.64 at 24 hours; Figure 2C). Additionally, Poly(I:C) stimulation at either time point led to the fewest induced pathways of all stimuli (125 up- and 1 downregulated at 4 hours, 241 up- and 17 downregulated at 24 hours; Figure D, Table S5 and S26). While we previously found 24 hour LPS-stimulated samples to be the most distinct from other samples stimulated with other pathogens at this time point (Figure 1B and S9B), we did not find a striking difference in the induced pathways for this stimulus compared to the other stimuli (Figure 2C and S11A, Table S5 and S26).

###### Differential expression analysis

We analyzed the genes differentially expressed in each of the four stimulus conditions at either time point, with RPMI medium as a control. We find that the 24 hour stimulation resulted in more DEGs than the 4 hour stimulation (mean N=1675.5 and N=379.8, Figure S13A). We find the DEGs at 4 hours include a higher fraction of known immune genes than the 24 hour DEGs (4hr up mean 41.3%, range 36.5-51.1%; 4hr down mean 27.2%, range 21.6-35.3%; 24hr up mean 34.4% range 31.3-37.9%; 24hr down mean 18.1%, range 14.2-27.8%; Figure S13A). Both the highest and lowest number of DEGs were found for 4 and 24 hour Poly(I:C) stimulation (N=2088 and N=45, Figure S13A, Table S8).

At the 4 hour time point, LPS and *C. albicans* show the most distinct DEGs (N=350, 40.1% and N=101, 11.6%), while these two conditions also share a high number of DEGs not found in the other conditions (N=133, 15.3%; Figure 3A, Table S8 and S28). We found 394 common upregulated DEGs at the 24 hour time point (11.9%; 326 upregulated, 82.7%; 68 downregulated, 17.3%). A high number of DEGs are specific to Poly(I:C) (N=636, 19.1%), overlapping between LPS, *C. albicans* and *S. aureus* (N=581, 17.5%), or specific to *S. aureus* stimulation (N=465, 14.0%, Figure 3B, Table S8 and S28), reflecting the pathogen clustering patterns identified previously (Figure 1C).

Core pathogen-stimulation DEGs

At the 4 hour time point, LPS and *C. albicans* show the most distinct DEGs (N=350, 40.1% and N=101, 11.6%), while these two conditions also share a high number of DEGs not found in the other conditions (N=133, 15.3%; Figure 3A, Table S8 and S28). We found 394 common upregulated DEGs at the 24 hour time point (11.9%; 326 upregulated, 82.7%; 68 downregulated, 17.3%). A high number of DEGs are specific to Poly(I:C) (N=636, 19.1%), overlapping between LPS, *C. albicans* and *S. aureus* (N=581, 17.5%), or specific to *S. aureus* stimulation (N=465, 14.0%, Figure 3B, Table S8 and S28), reflecting the pathogen clustering patterns identified previously (Figure 1C).
