## Supplementary figures for "Transcription-based identification of uncharacterized genes in the human immune response"

### Slide 1
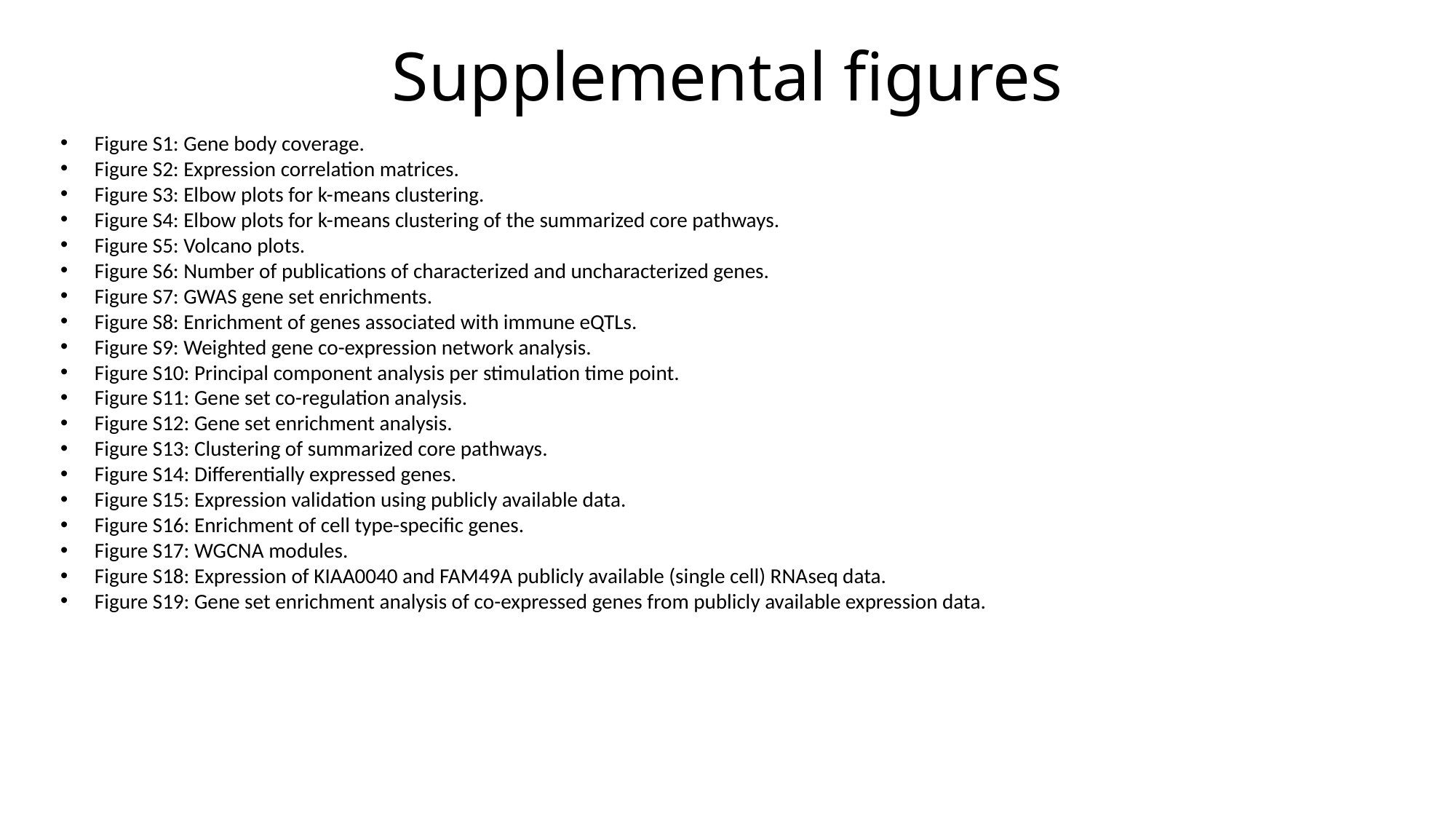

#### Supplemental figures
Figure S1: Gene body coverage.
Figure S2: Expression correlation matrices.
Figure S3: Elbow plots for k-means clustering.
Figure S4: Elbow plots for k-means clustering of the summarized core pathways.
Figure S5: Volcano plots.
Figure S6: Number of publications of characterized and uncharacterized genes.
Figure S7: GWAS gene set enrichments.
Figure S8: Enrichment of genes associated with immune eQTLs.
Figure S9: Weighted gene co-expression network analysis.
Figure S10: Principal component analysis per stimulation time point.
Figure S11: Gene set co-regulation analysis.
Figure S12: Gene set enrichment analysis.
Figure S13: Clustering of summarized core pathways.
Figure S14: Differentially expressed genes.
Figure S15: Expression validation using publicly available data.
Figure S16: Enrichment of cell type-specific genes.
Figure S17: WGCNA modules.
Figure S18: Expression of KIAA0040 and FAM49A publicly available (single cell) RNAseq data.
Figure S19: Gene set enrichment analysis of co-expressed genes from publicly available expression data.

### Slide 2
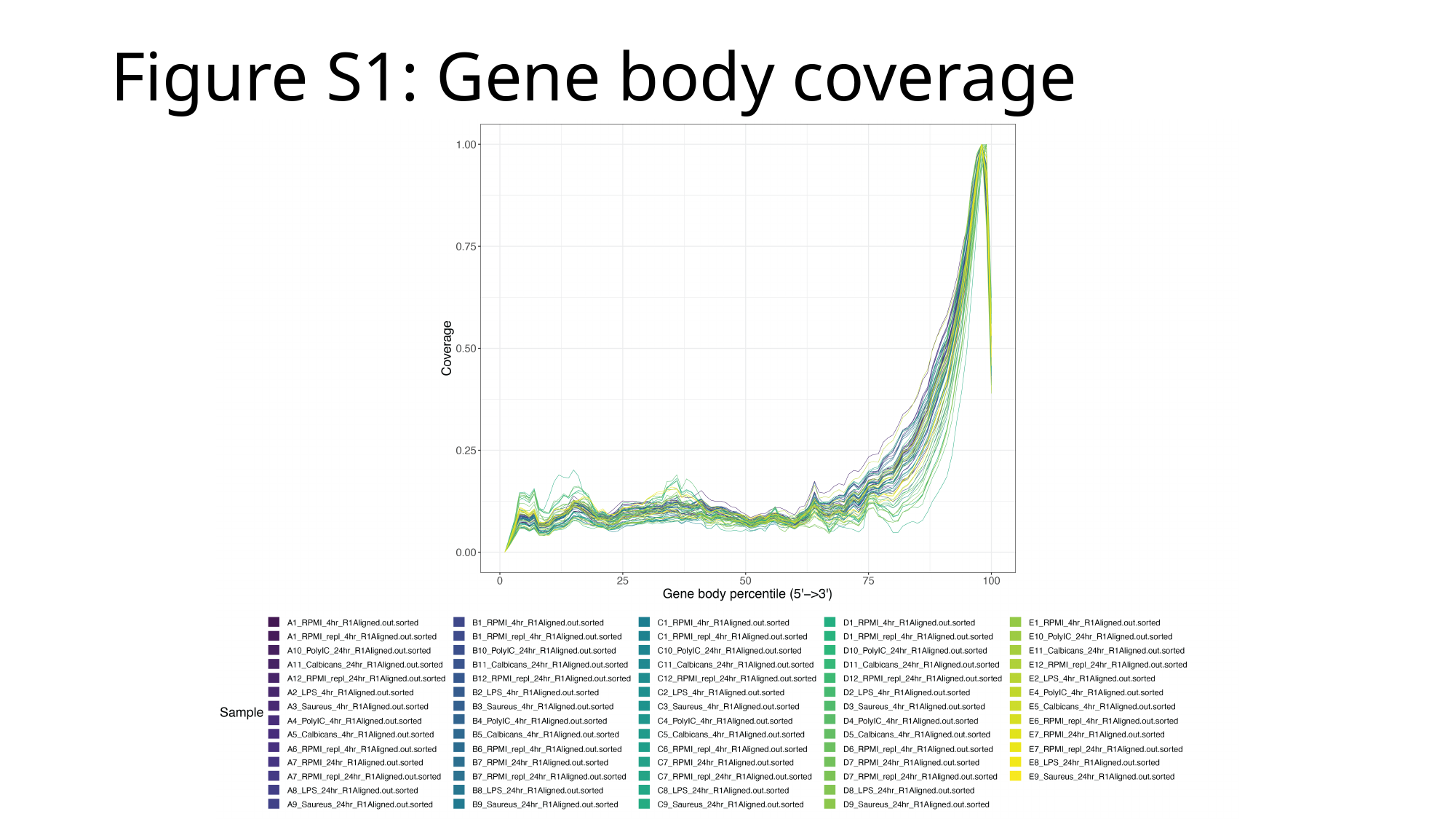

#### Figure S1: Gene body coverage

### Slide 3
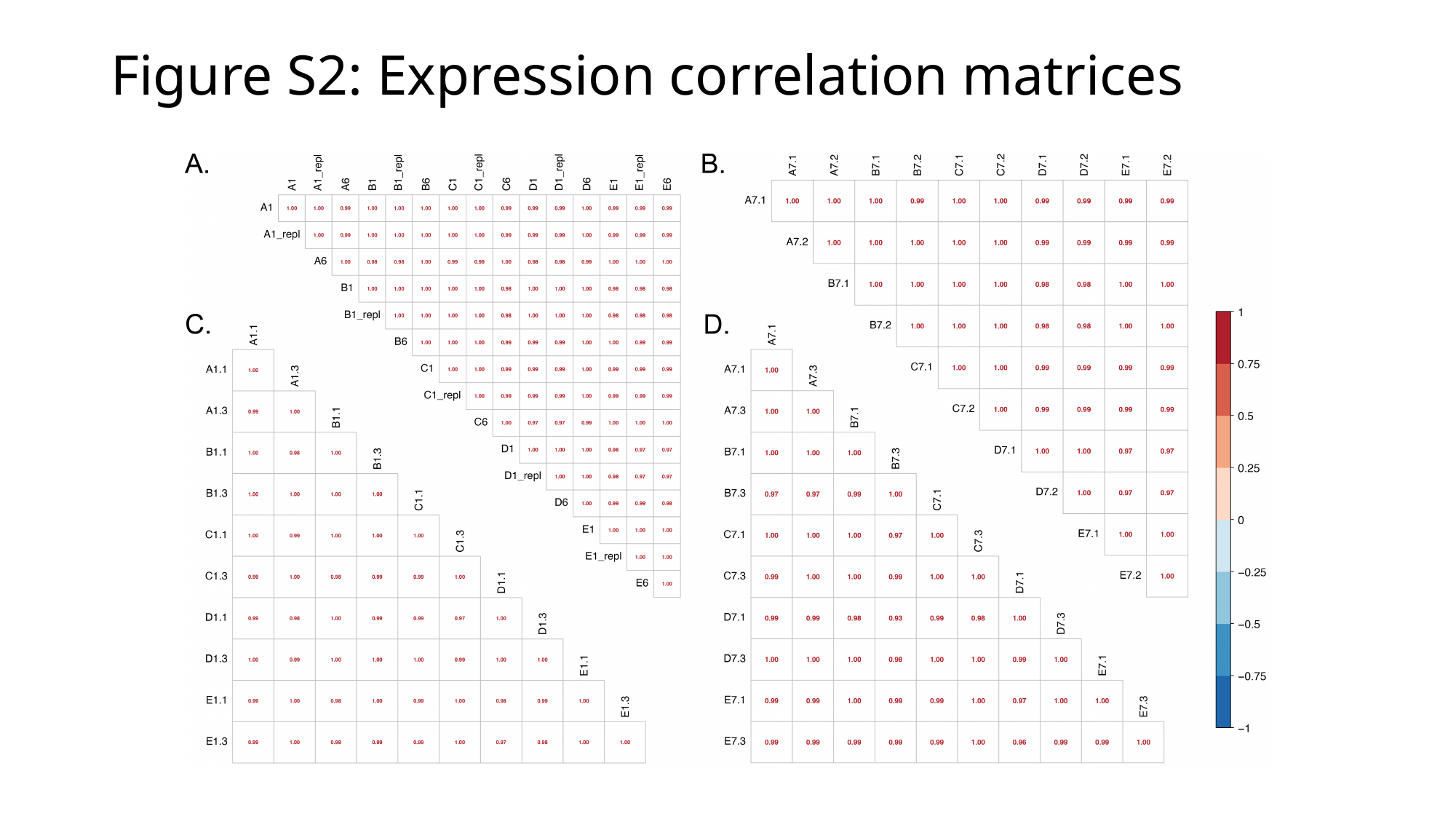

#### Figure S2: Expression correlation matrices

### Slide 4
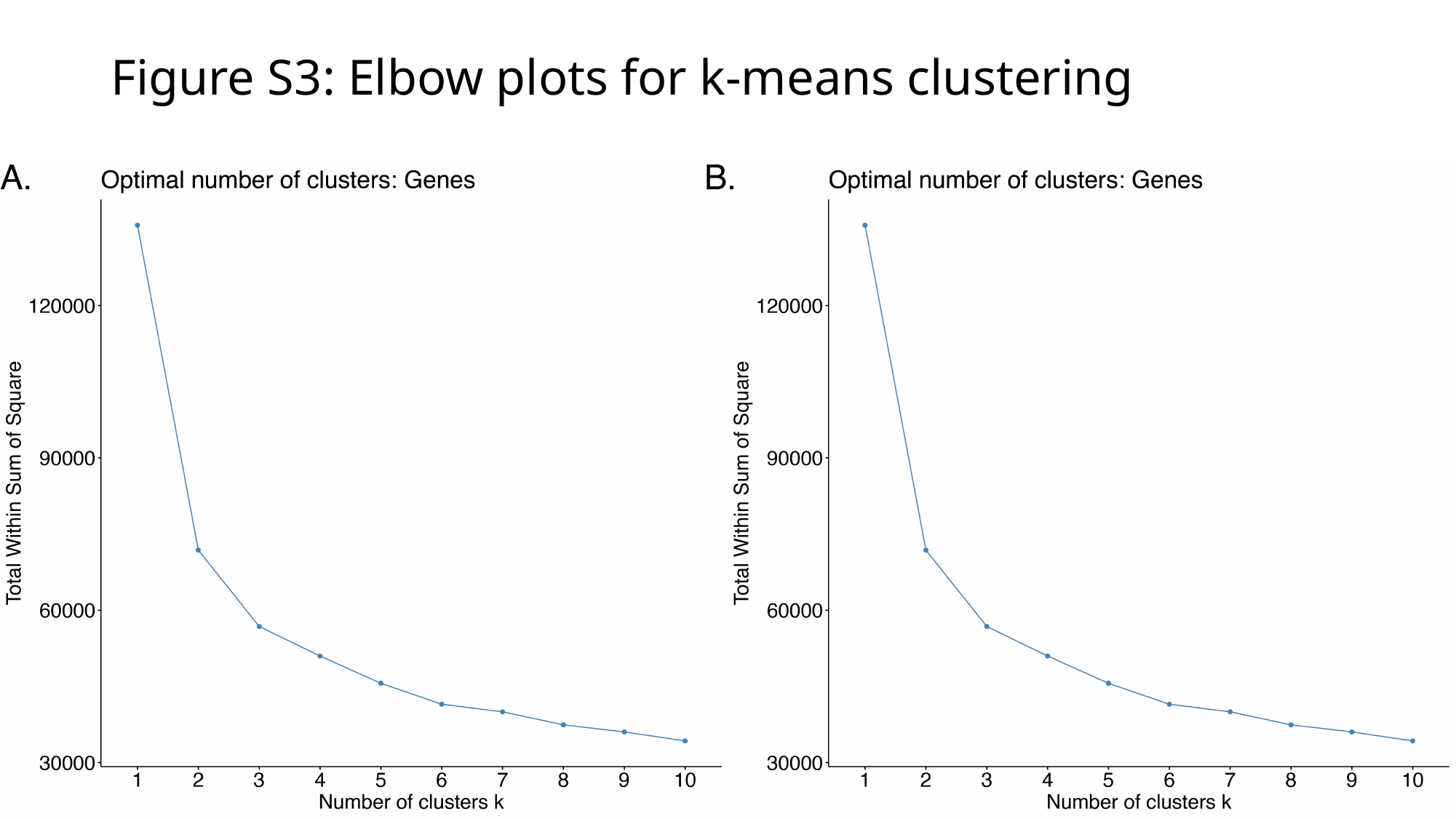

#### Figure S3: Elbow plots for k-means clustering

### Slide 5
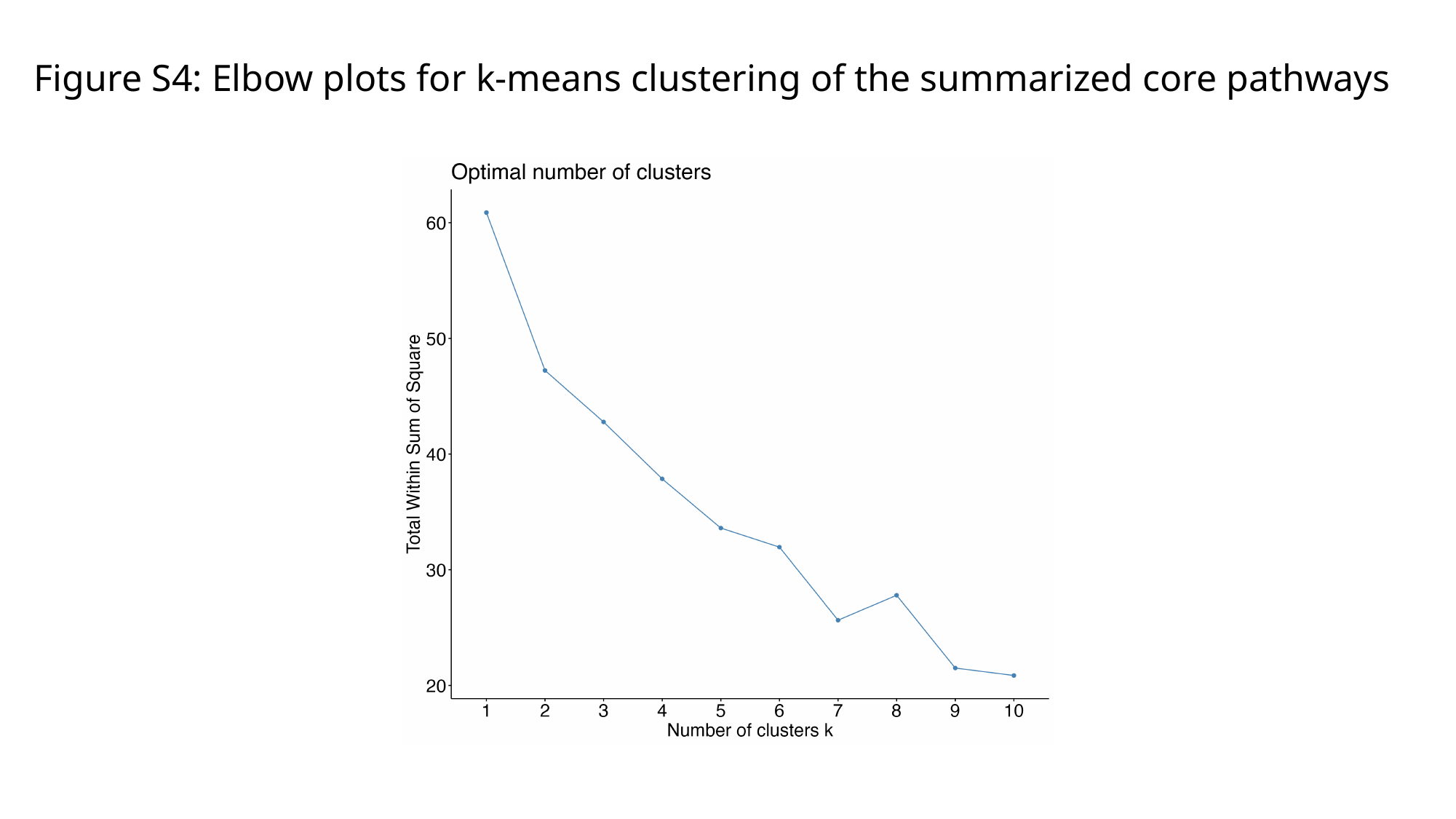

#### Figure S4: Elbow plots for k-means clustering of the summarized core pathways

### Slide 6
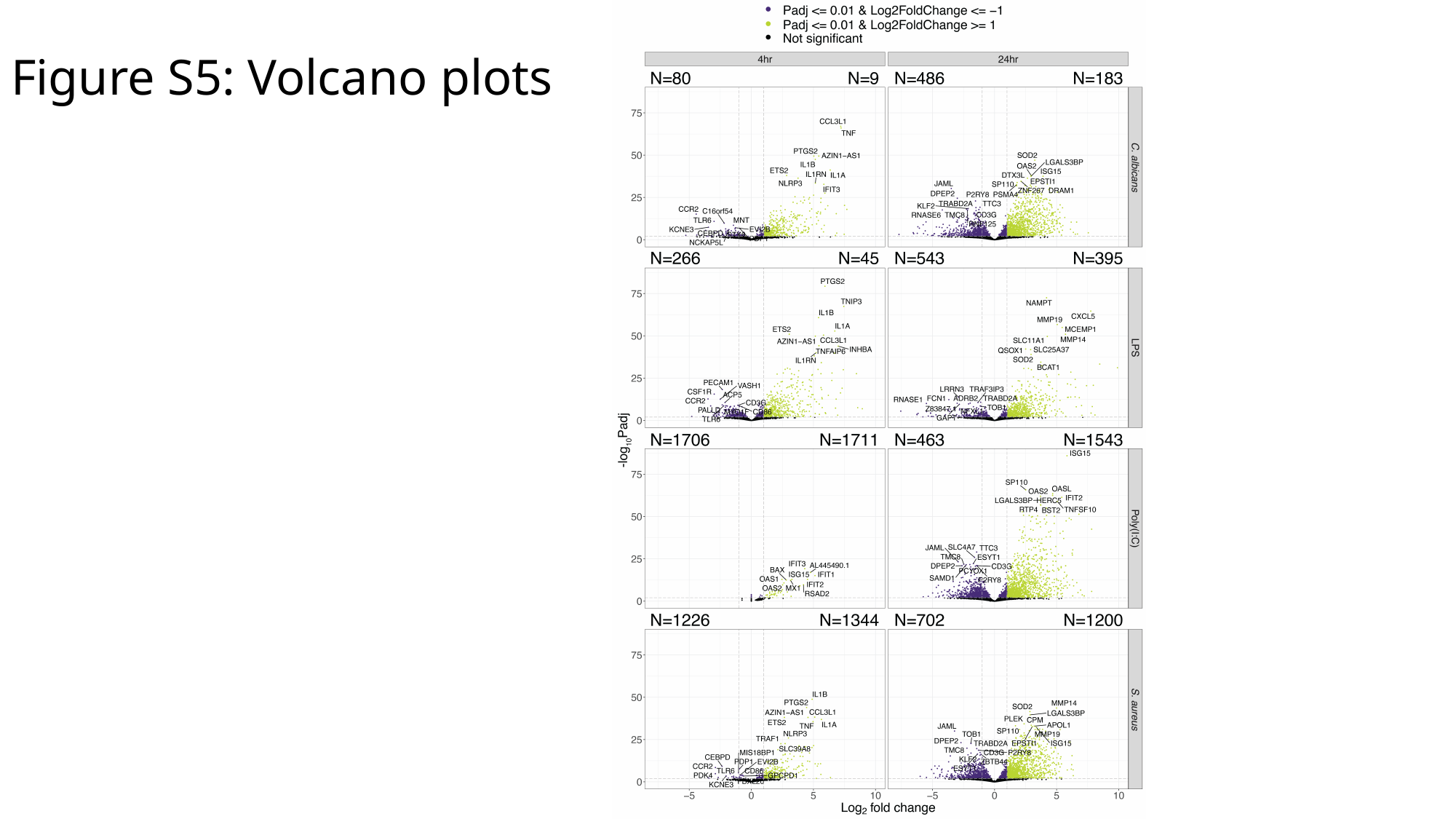

#### Figure S5: Volcano plots

### Slide 7
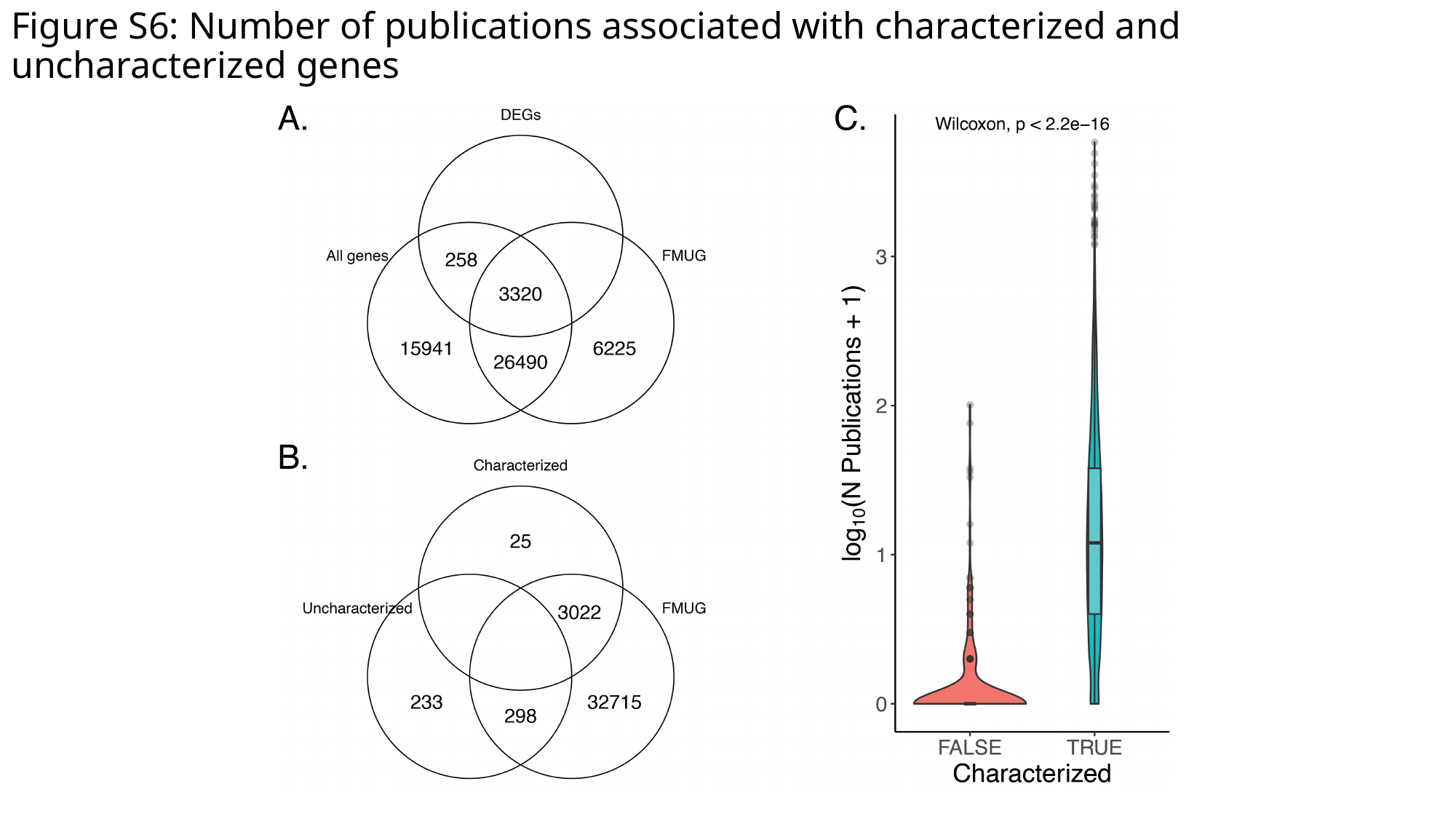

#### Figure S6: Number of publications associated with characterized and uncharacterized genes

### Slide 8
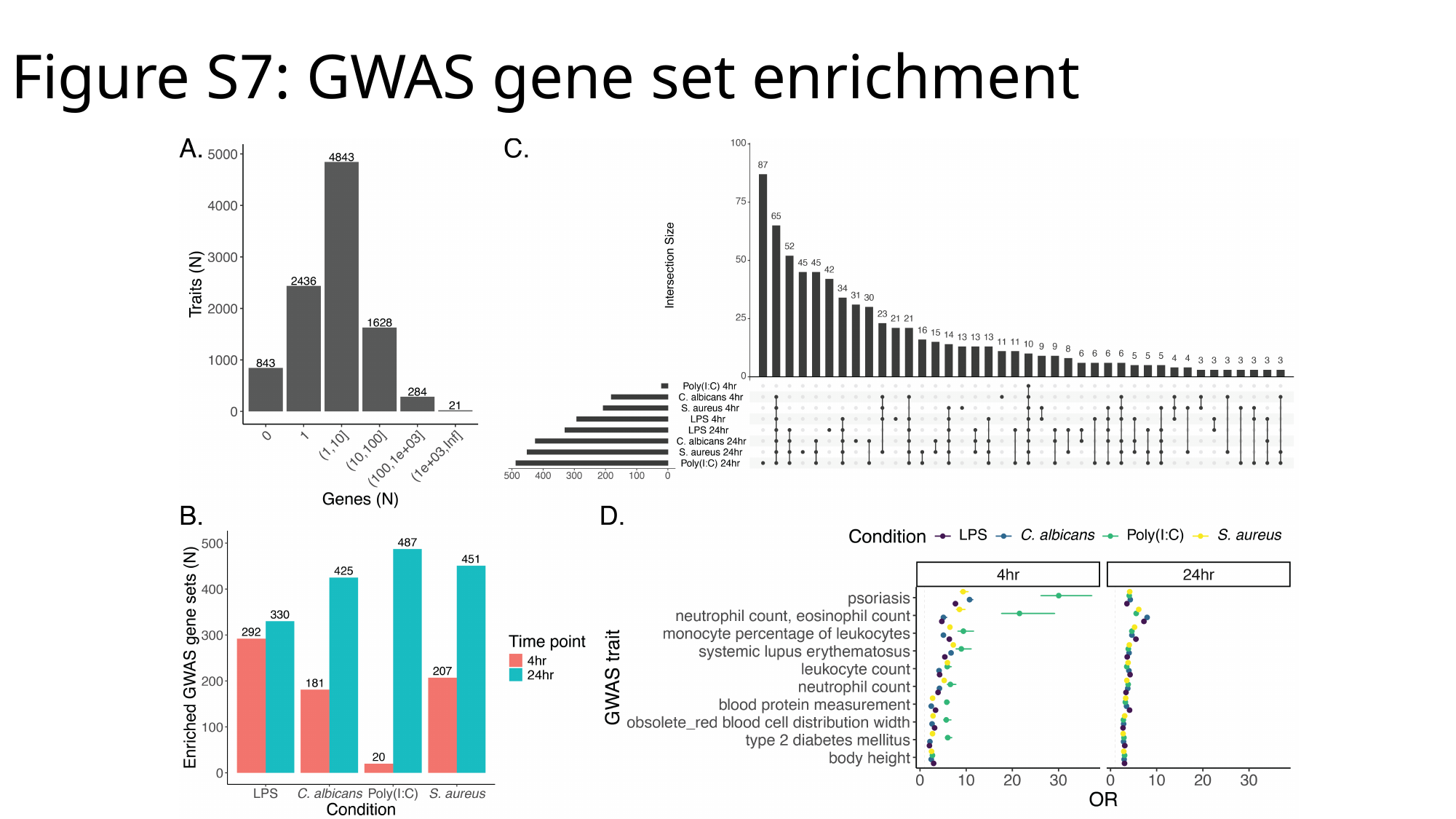

#### Figure S7: GWAS gene set enrichment

### Slide 9
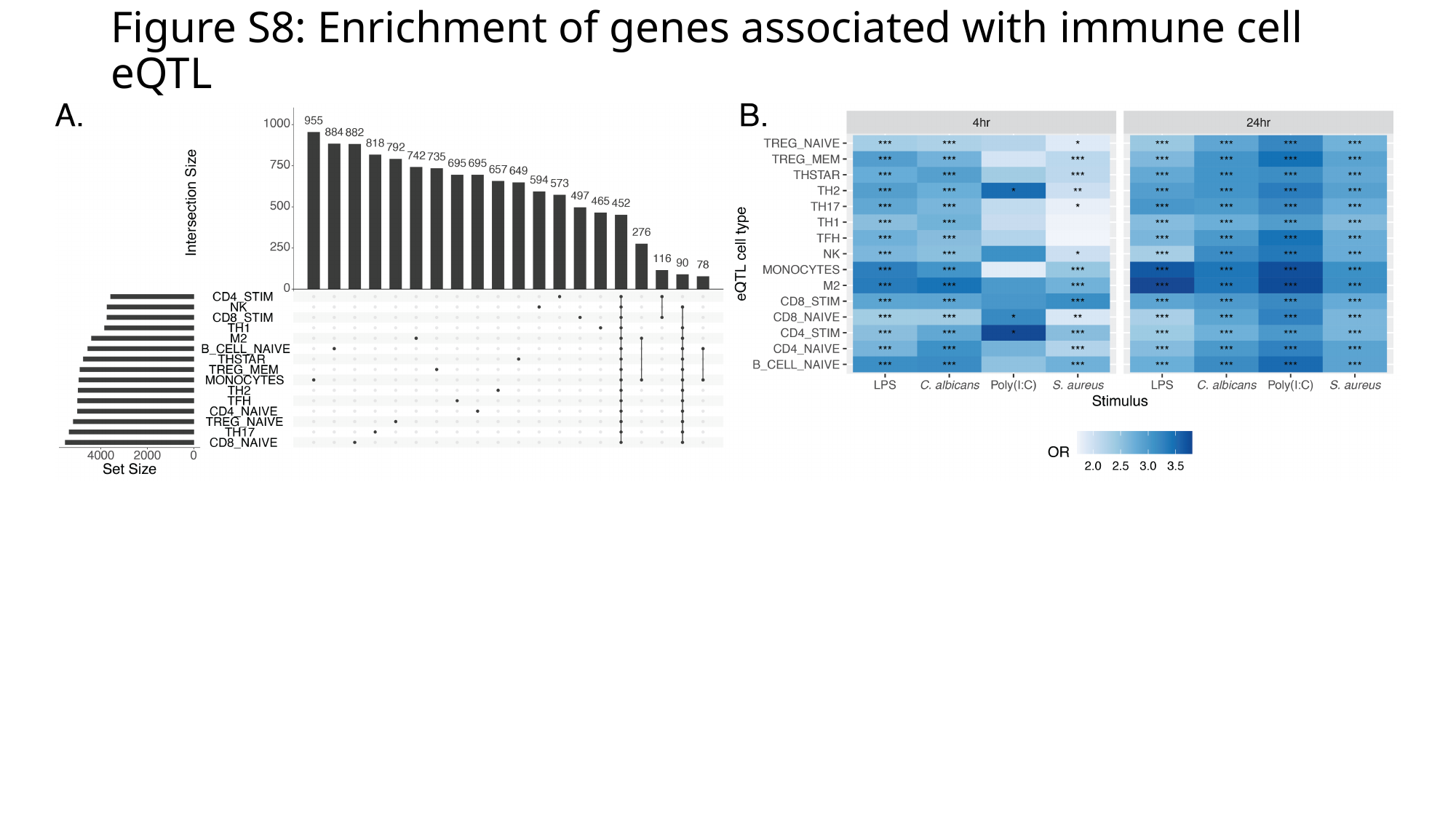

#### Figure S8: Enrichment of genes associated with immune cell eQTL

### Slide 10
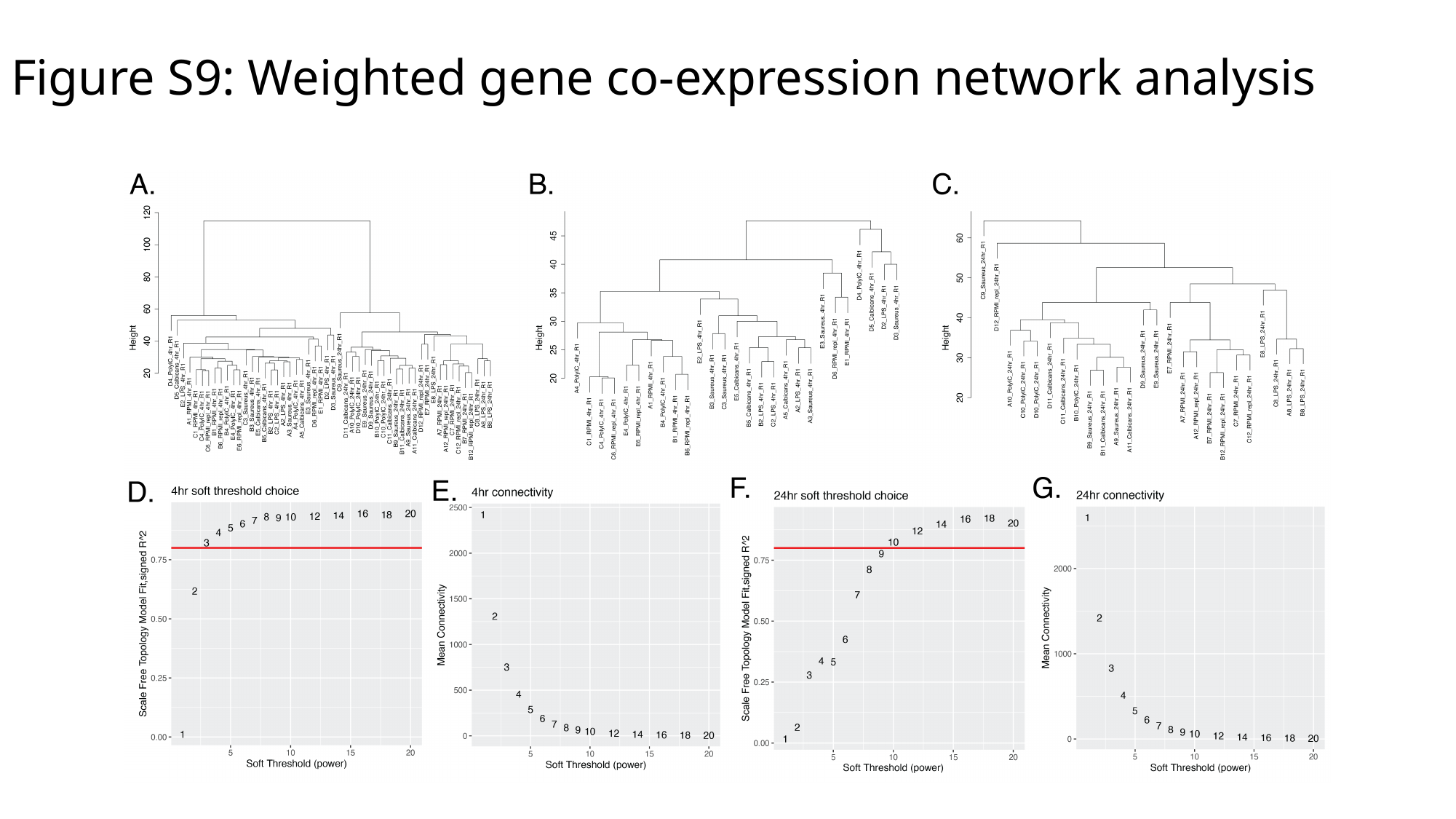

#### Figure S9: Weighted gene co-expression network analysis

### Slide 11
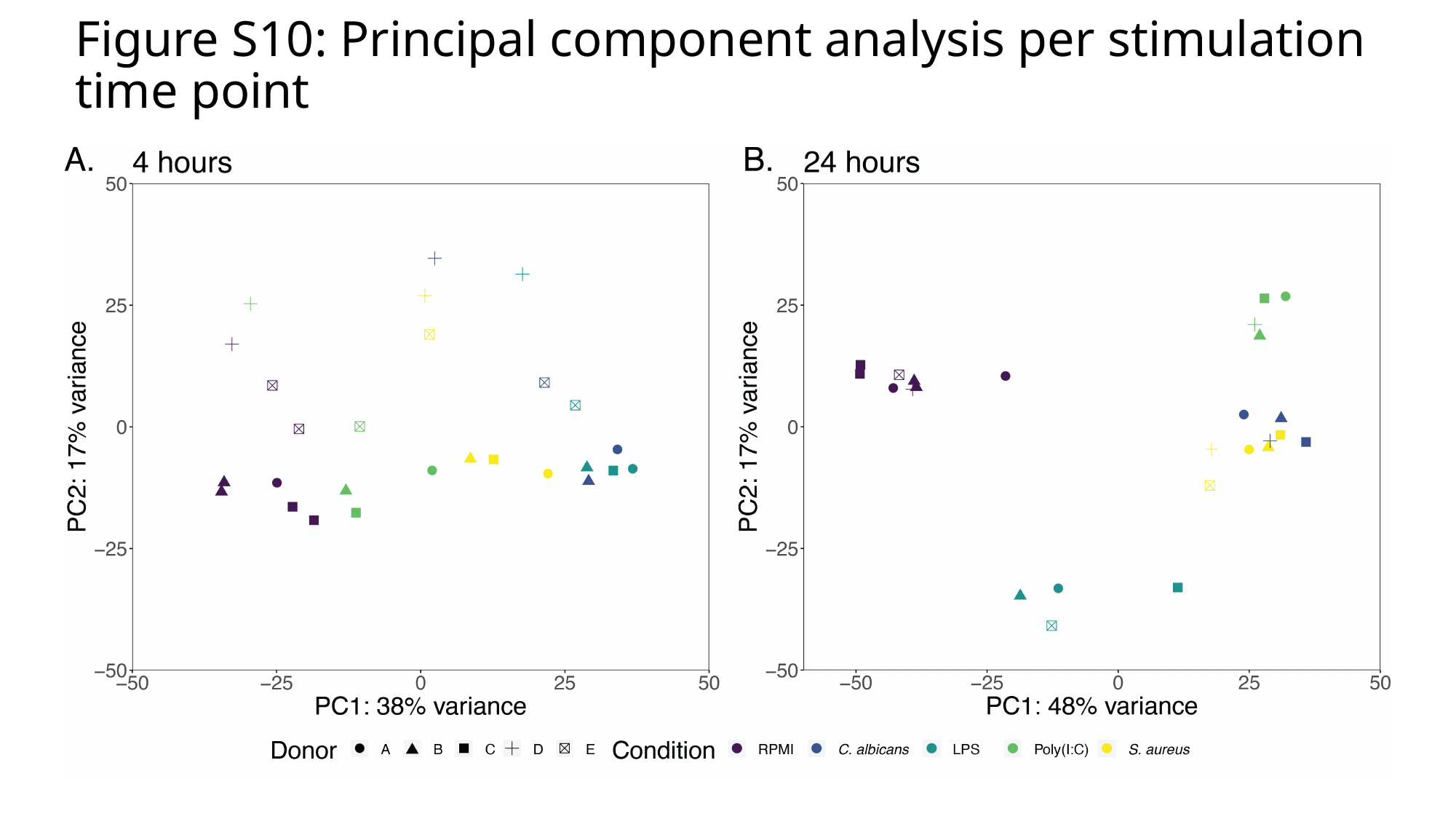

#### Figure S10: Principal component analysis per stimulation time point

### Slide 12
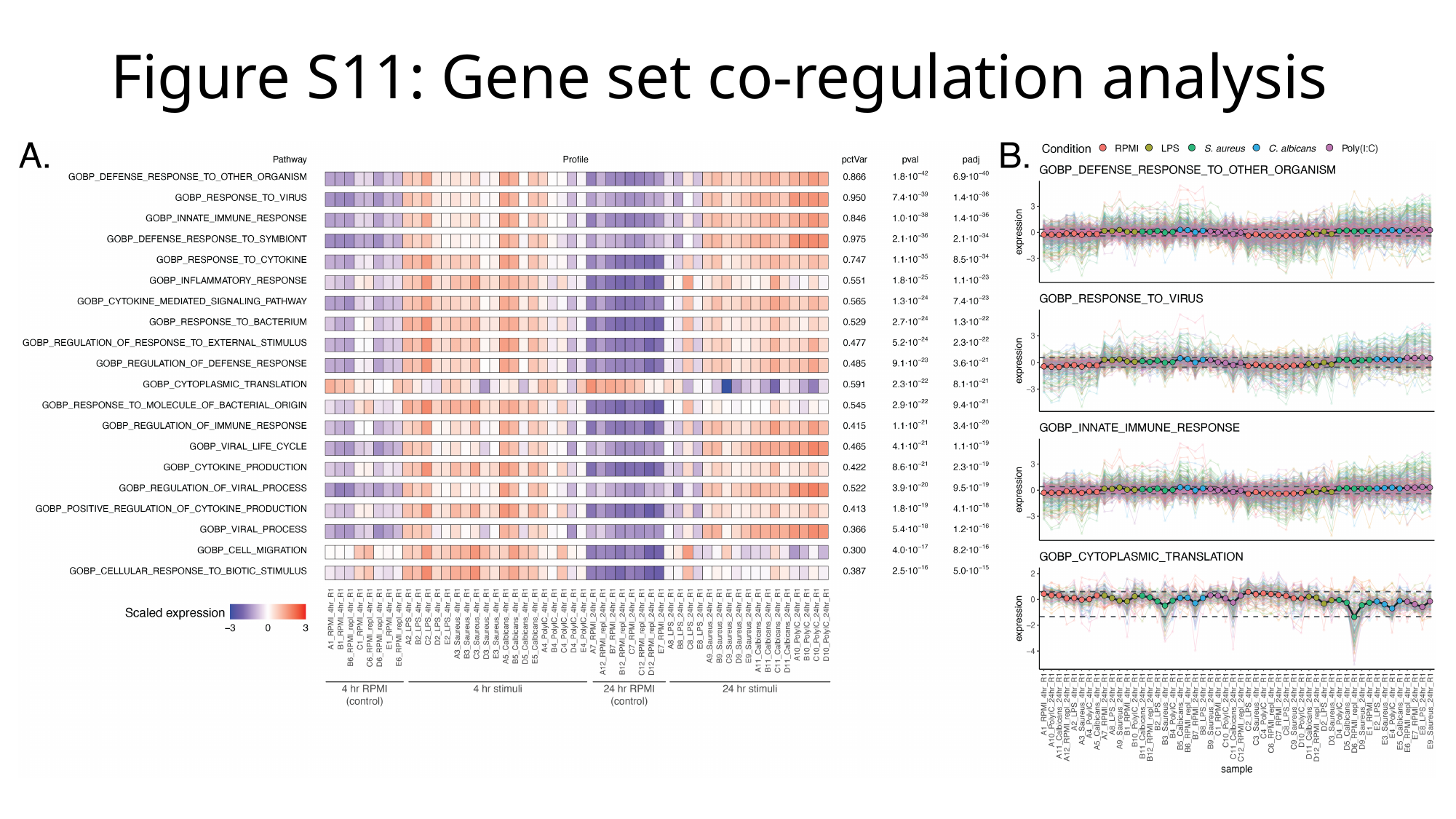

#### Figure S11: Gene set co-regulation analysis

### Slide 13
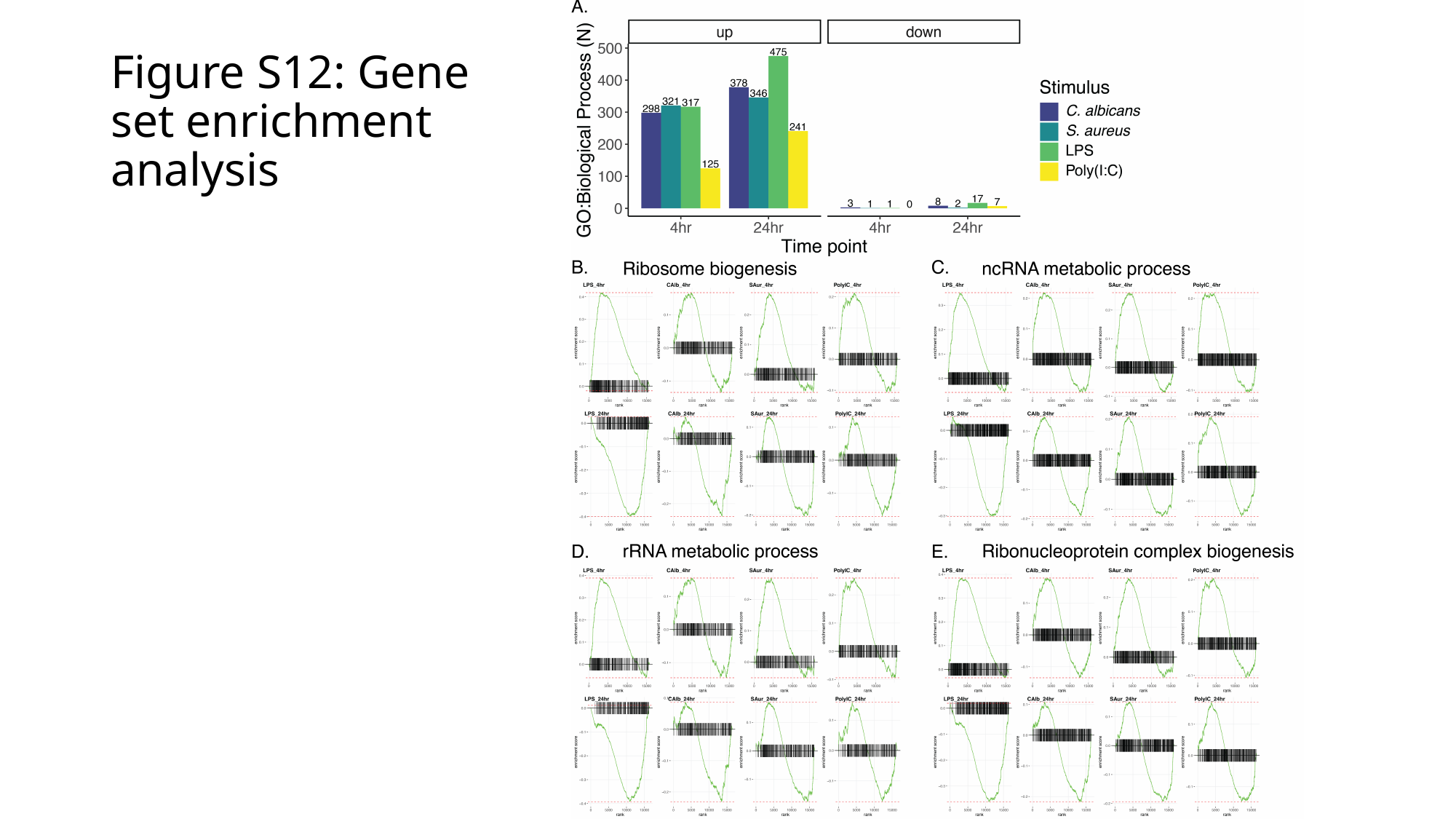

#### Figure S12: Gene set enrichment analysis

### Slide 14
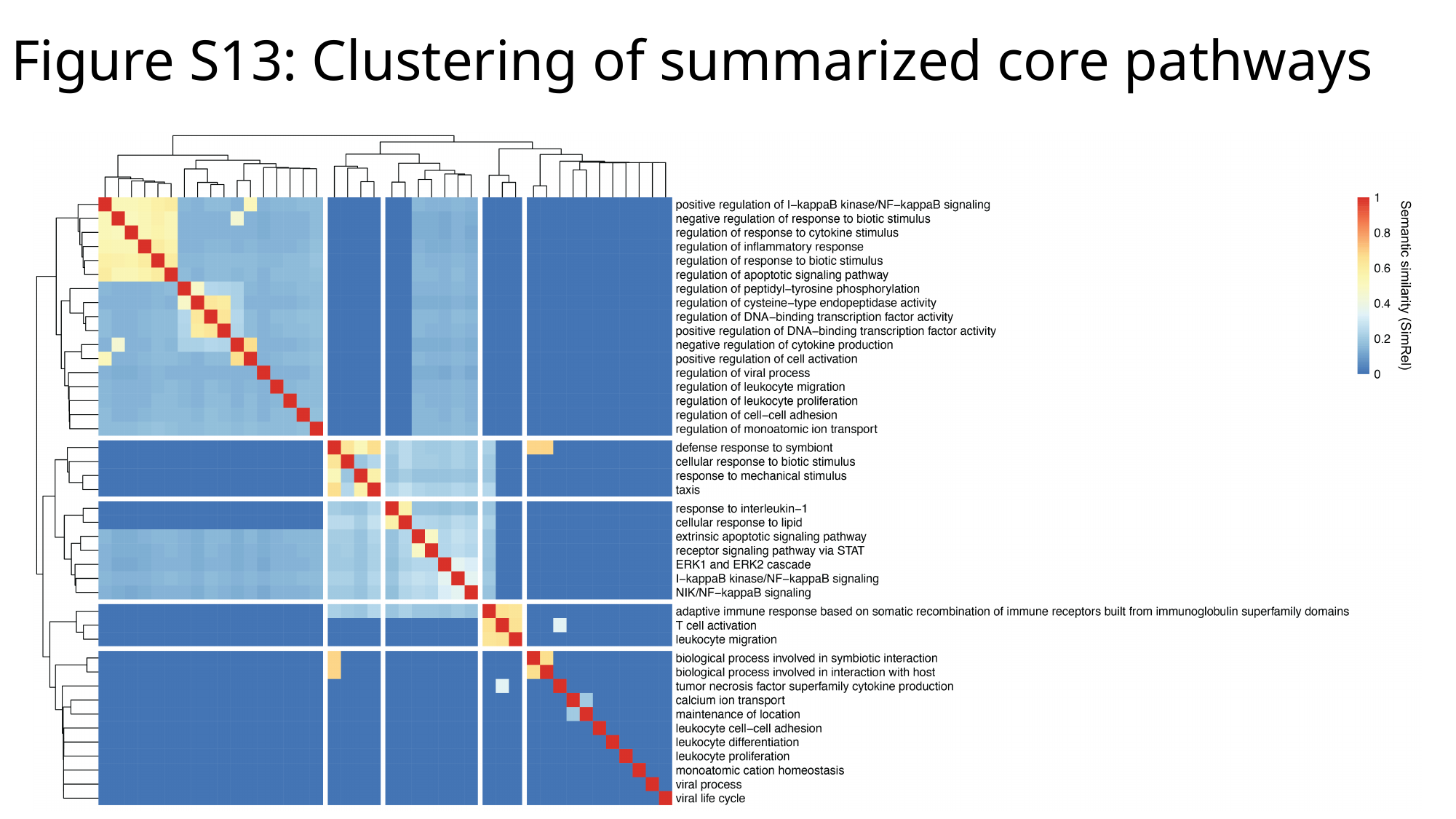

#### Figure S13: Clustering of summarized core pathways

### Slide 15
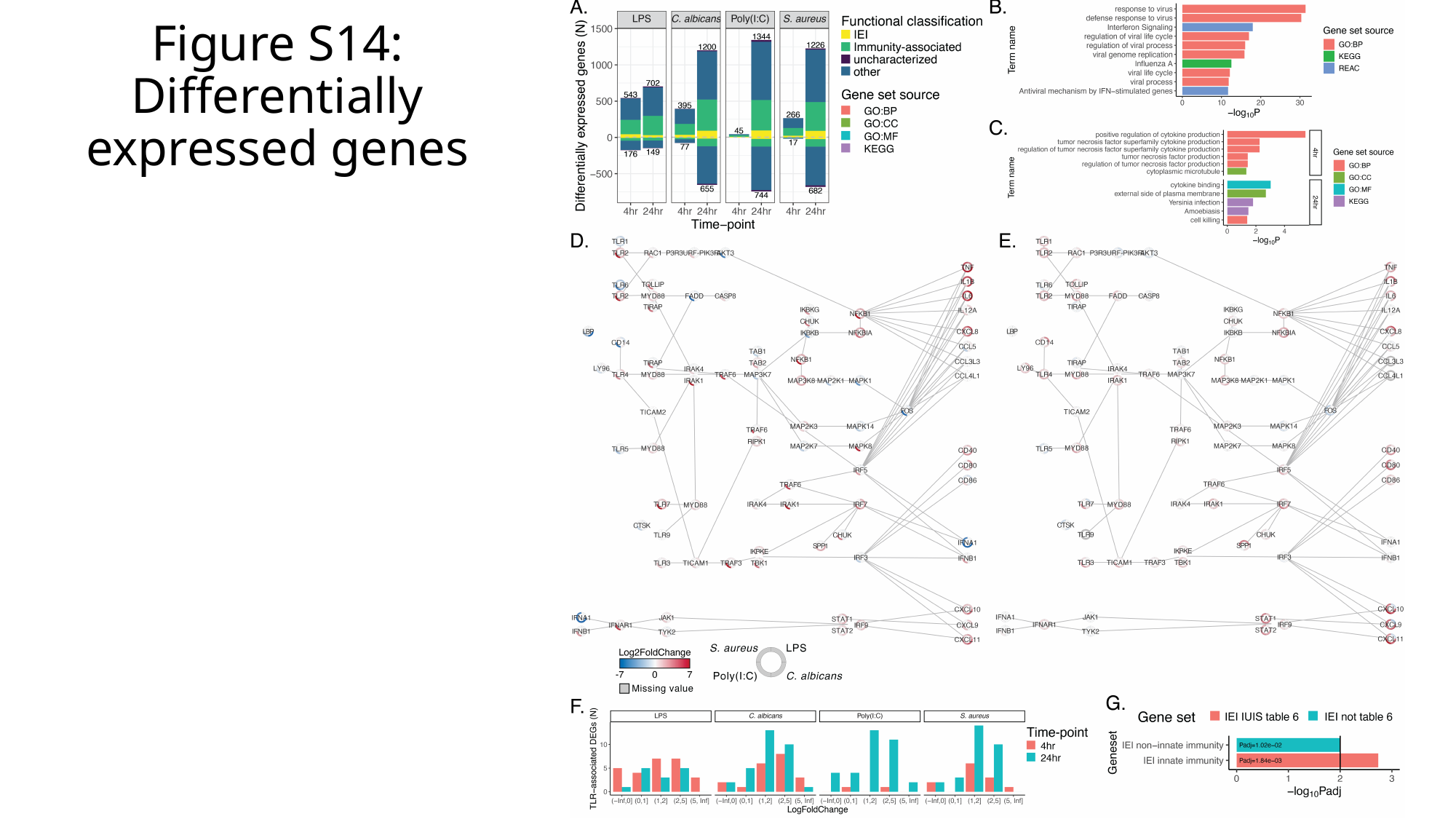

#### Figure S14: Differentially expressed genes

### Slide 16
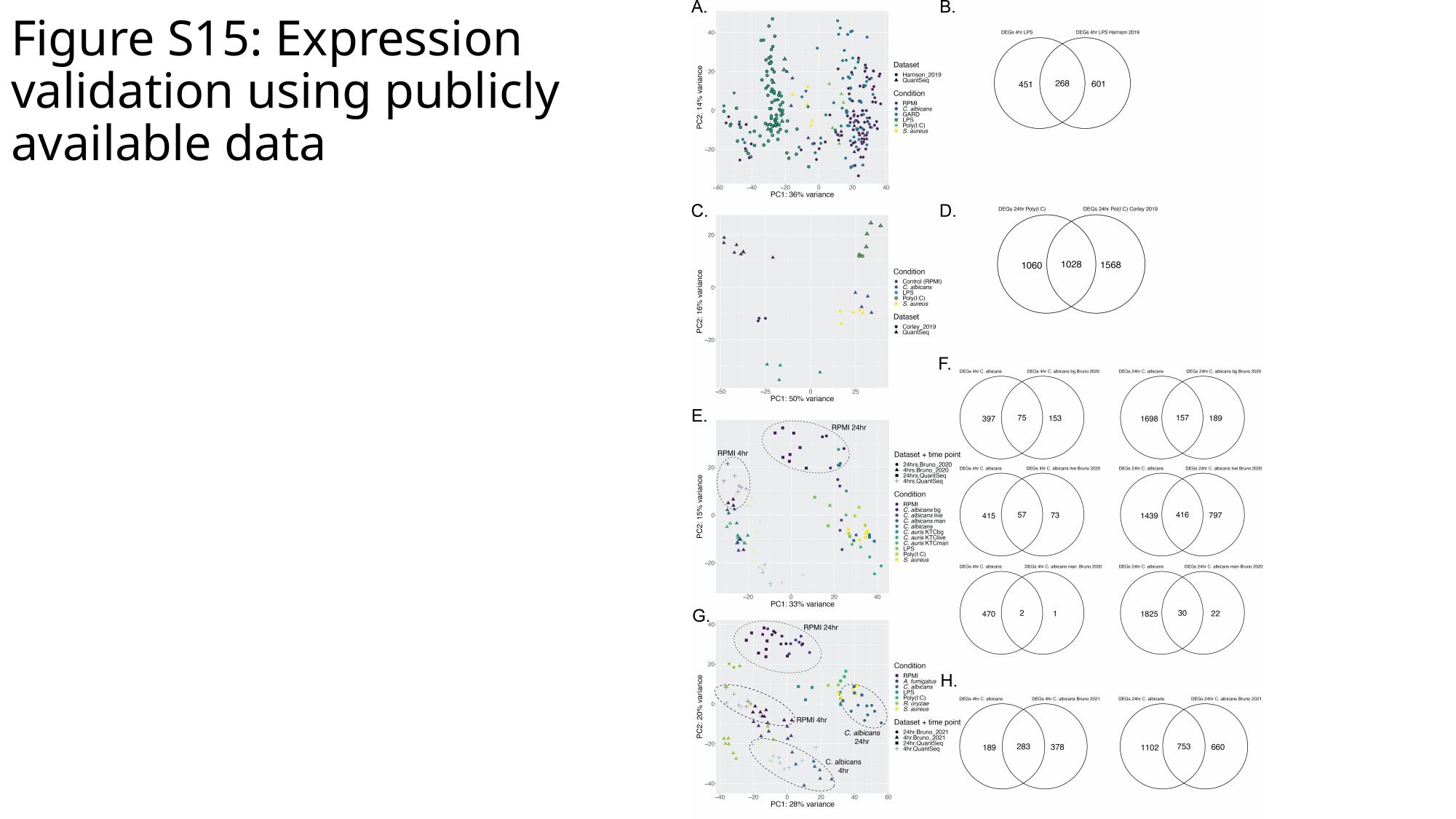

#### Figure S15: Expression validation using publicly available data

### Slide 17
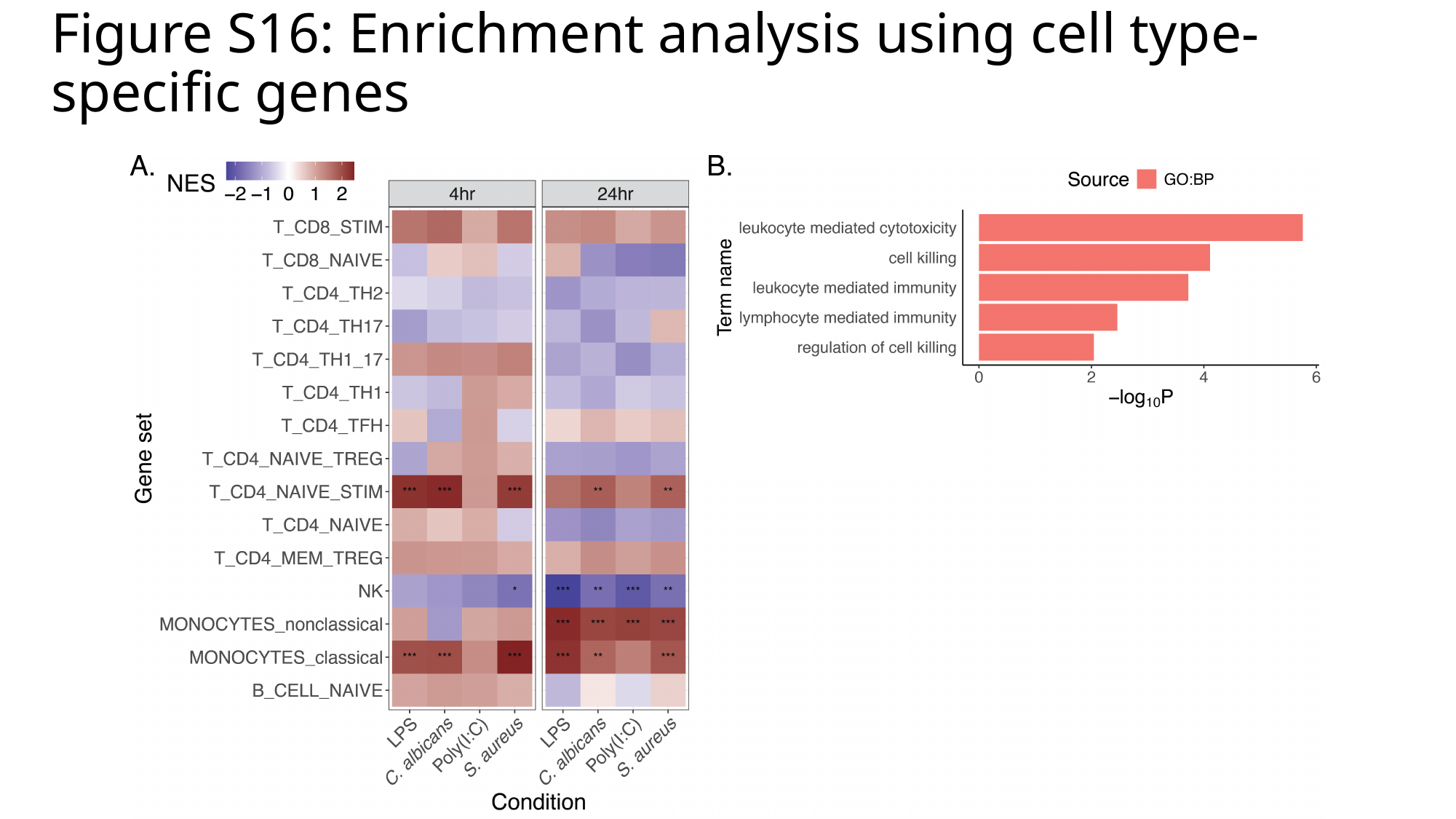

#### Figure S16: Enrichment analysis using cell type-specific genes

### Slide 18
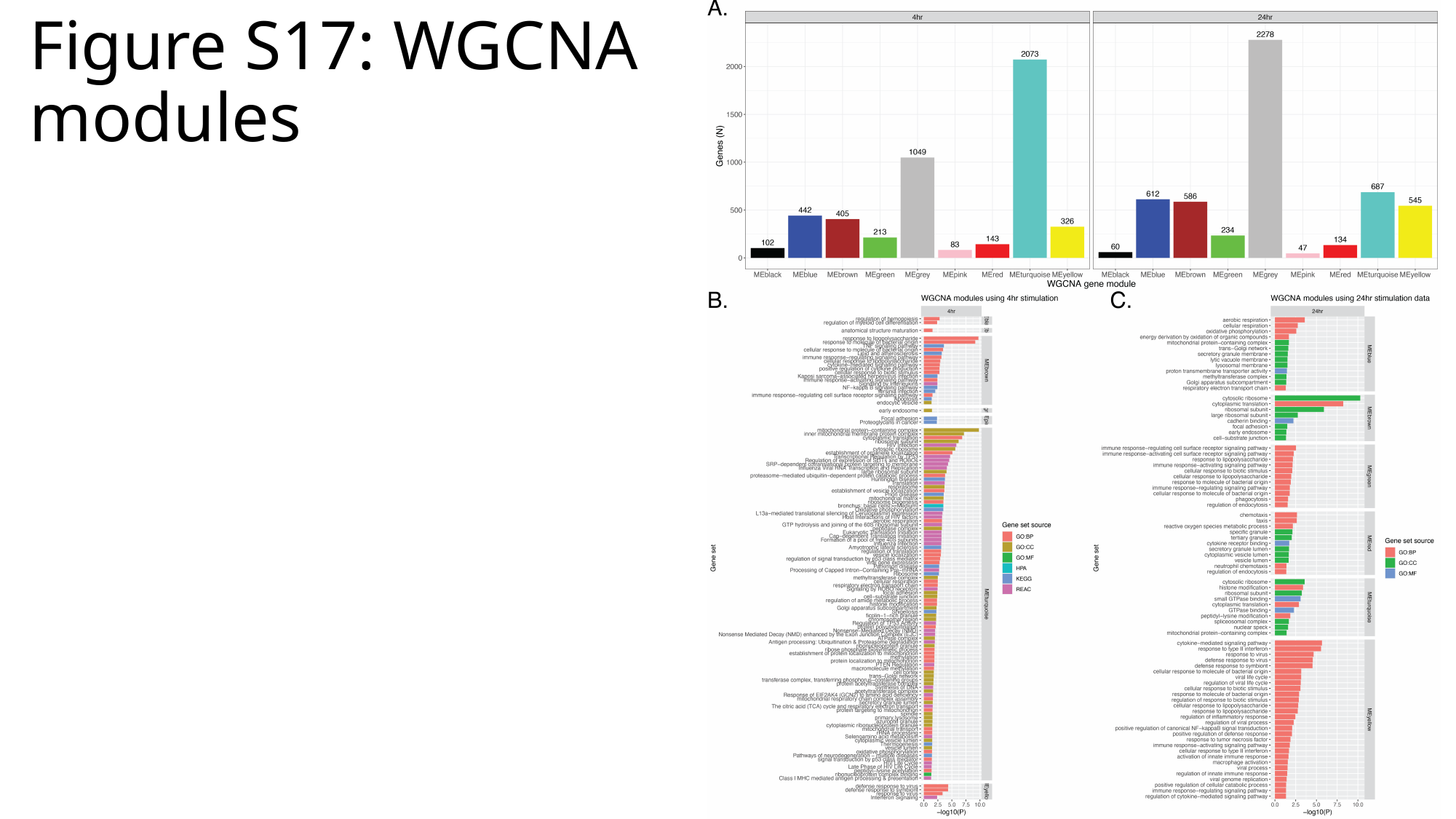

#### Figure S17: WGCNA modules

### Slide 19
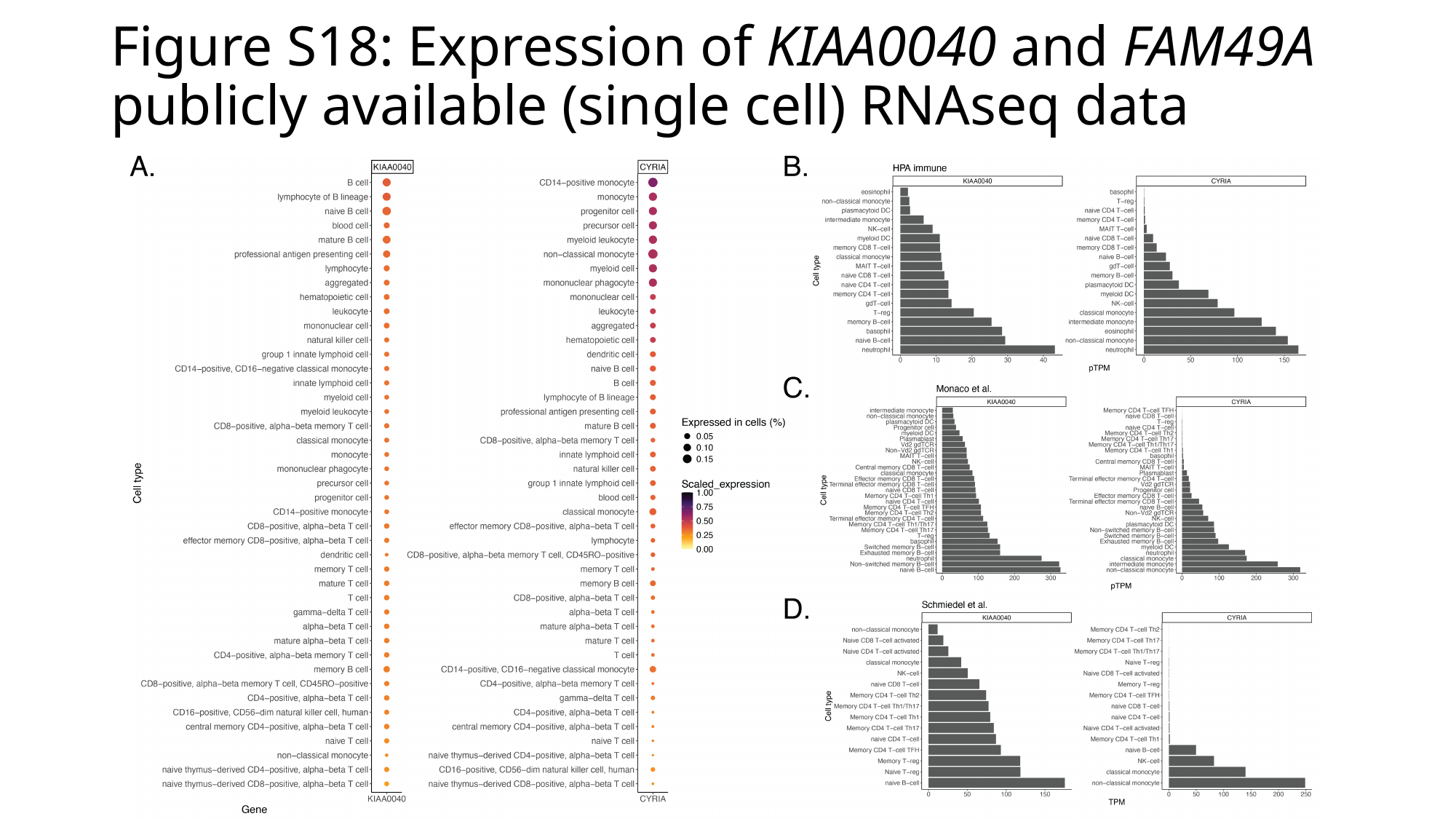

#### Figure S18: Expression of KIAA0040 and FAM49A publicly available (single cell) RNAseq data

### Slide 20
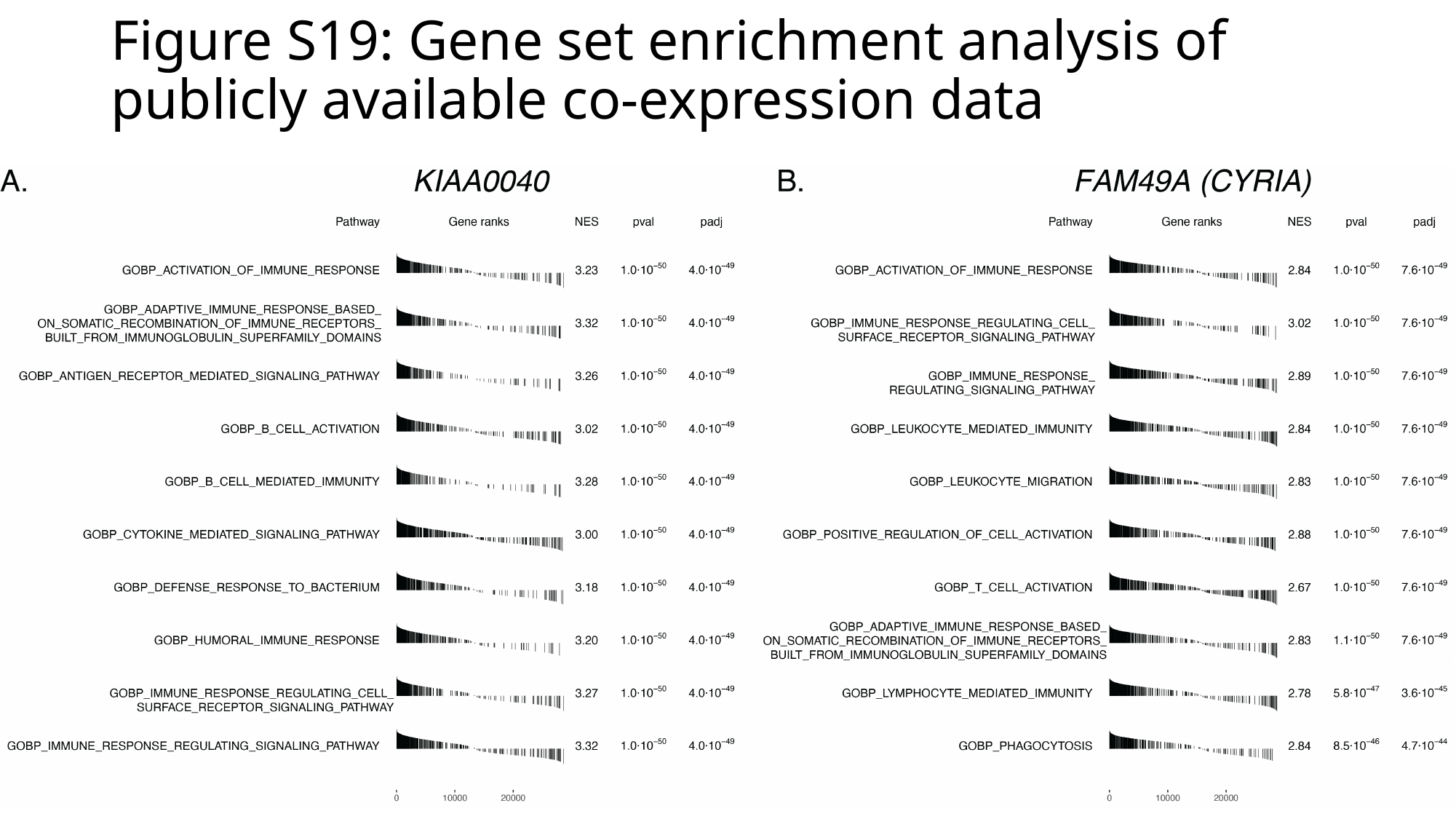

#### Figure S19: Gene set enrichment analysis of publicly available co-expression data
